# Lactate activates the mitochondrial electron transport chain independent of its metabolism

**DOI:** 10.1101/2023.08.02.551712

**Authors:** Xin Cai, Charles C. Ng, Olivia Jones, Tak Shun Fung, Keunwoo Ryu, Dayi Li, Craig B. Thompson

**Affiliations:** Cancer Biology and Genetics Program, Memorial Sloan Kettering Cancer Center, New York, NY 10065, USA; Department of Radiation Oncology, Memorial Sloan Kettering Cancer Center, New York, NY 10065, USA

**Keywords:** lactate, mitochondria, oxidative phosphorylation, TCA cycle, electron transport chain, glycolysis.

## Abstract

Lactate has long been considered a cellular waste product. However, we found that as extracellular lactate accumulates, it also enters the mitochondrial matrix and stimulates mitochondrial electron transport chain (ETC) activity. The resulting increase in mitochondrial ATP synthesis suppresses glycolysis and increases the utilization of pyruvate and/or alternative respiratory substrates. The ability of lactate to increase oxidative phosphorylation does not depend on its metabolism. Both L- and D-lactate are effective at enhancing ETC activity and suppressing glycolysis. Furthermore, the selective induction of mitochondrial oxidative phosphorylation by unmetabolized D-lactate reversibly suppressed aerobic glycolysis in both cancer cell lines and proliferating primary cells in an ATP-dependent manner and enabled cell growth on respiratory-dependent bioenergetic substrates. In primary T cells, D-lactate enhanced cell proliferation and effector function. Together, these findings demonstrate that lactate is a critical regulator of the ability of mitochondrial oxidative phosphorylation to suppress glucose fermentation.

## INTRODUCTION

As a glycolytic byproduct, lactate is rapidly produced during periods of intense exercise and increased ATP demand. This led to the common belief that lactate is a metabolic waste product. However, lactate can be used as a fuel for the mitochondrial TCA cycle during wound repair and tissue regeneration^1–4^. More recent in vivo isotope tracing studies have indicated that extracellular lactate, instead of glucose, is used as the primary substrate to support the TCA cycle in most major organs and tumors under basal metabolic conditions^5–8^. However, the mechanism by which lactate serves as the preferential carbon source for the mitochondria remains unknown. These observations raise the question of how cells sense the availability of lactate as a nutrient source. Furthermore, lactate has been shown to reprogram cancer and immune cell function through unclear mechanisms^9–11^, suggesting lactate has additional metabolic regulatory functions in addition to being an oxidizable substrate.

At 1-2 mM physiological concentration, lactate is the second most abundant circulating carbon source after glucose and is produced primarily through glycolysis. Since lactate is not appreciably excreted from the body, circulating lactate is either oxidized to CO2 in the mitochondria to produce ATP or used to build carbon backbones of macromolecules. The known metabolic functions of lactate are dependent on the cytosolic lactate dehydrogenase (LDH) enzyme that converts lactate to pyruvate while reducing NAD^+^ into NADH. The resulting pyruvate can then be imported into the mitochondria and be used to support TCA cycle-dependent oxidative phosphorylation. The electrons from pyruvate oxidation are ultimately deposited into the mitochondrial electron transport chain (ETC) for ATP production. For lactate to serve as a meaningful mitochondrial energy source therefore requires its LDH-dependent conversion to pyruvate and pyruvate entry into the TCA cycle through the pyruvate dehydrogenase (PDH) complex. Whether there are LDH-independent metabolic regulatory functions of lactate is unclear.

Here, we show that lactate activates the ETC to increase mitochondrial ATP production independent of its metabolism. The ability of lactate to stimulate the ETC is independent of LDH and also does not depend on pyruvate entry into the mitochondria or the TCA cycle but requires oxygen availability. Lactate-induced increase in oxidative phosphorylation results in more pyruvate entry into the TCA cycle, thus further increasing mitochondrial ATP production while suppressing glycolysis. Therefore, lactate serves as a mitochondrial messenger to shift ATP production from glycolysis to oxidative phosphorylation, allowing cells to conserve glucose while using lactate-derived pyruvate as their preferential substrate to support cellular ATP production.

## RESULTS

### Lactate supports cell growth under limiting glucose and suppresses glycolysis

To first investigate lactate’s function as a carbon source, we asked whether it could replace glucose as an anabolic substrate. When HepG2 cells were cultured in complete tissue culture medium containing 10 mM glucose, the cells engaged in sustained proliferation, expanding over tenfold in a week (Figure 1A). However, when the glucose in the medium was replaced with an equivalent amount of reduced carbon in the form of L-lactate (20 mM), cells were unable to maintain their viability and proliferation. We next reduced the level of glucose until it became limiting for growth^12^. Under limiting glucose (1mM glucose), the addition of 20 mM L-lactate substantially restored cell proliferation while unexpectedly reducing glucose consumption (Figures 1A and 1B). The ability of lactate to suppress glycolysis was universally observed across glucose concentrations and multiple cell lines (Figures S1A and S1B).

**Figure 1.**
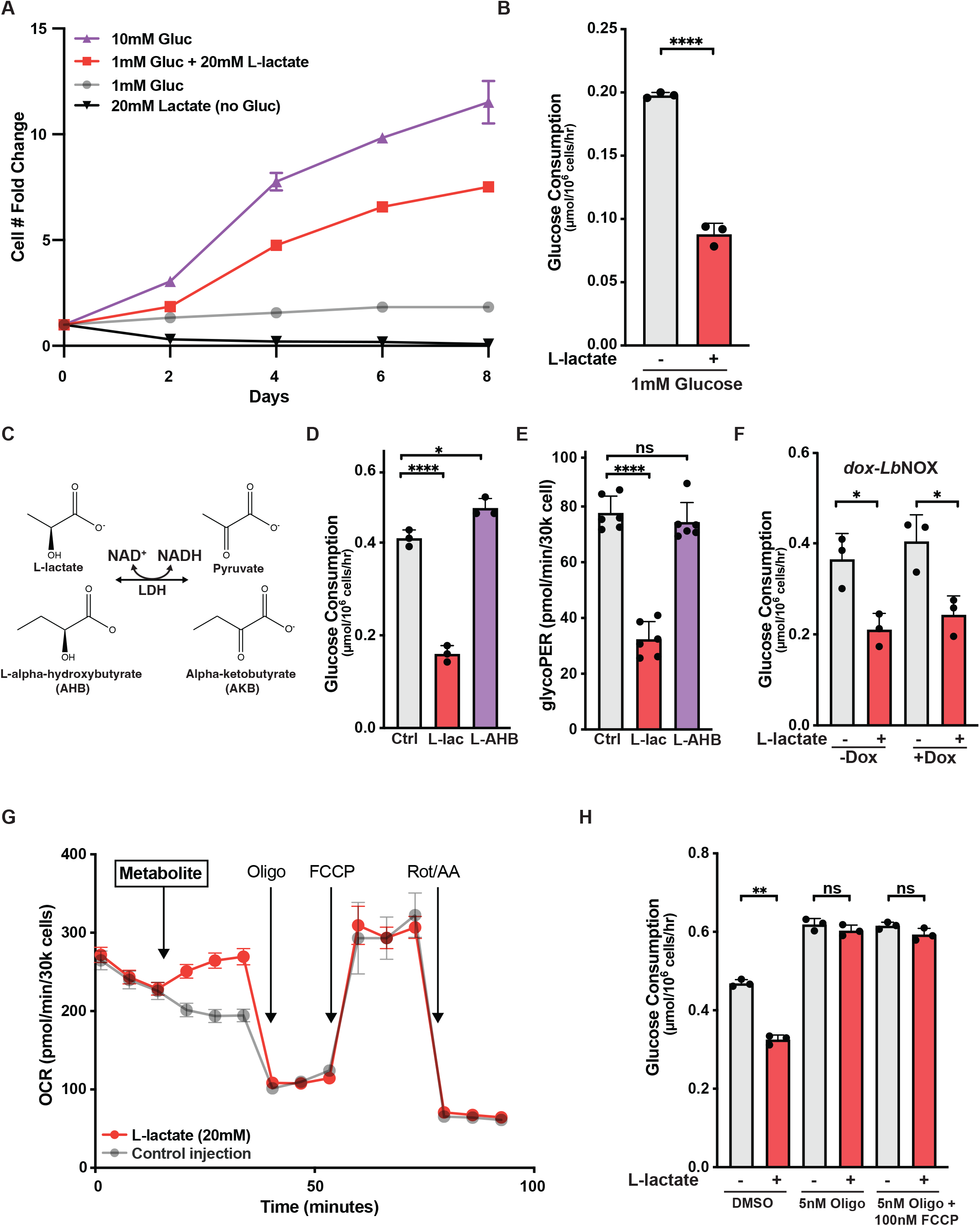
Lactate suppresses glycolysis by increasing oxidative phosphorylation. (A) Proliferation of HepG2 cells cultured in the indicated glucose and lactate concentrations measured as cell number fold change relative to day 0. Gluc, glucose. (B) Glucose consumption of HepG2 cells cultured in medium with the indicated glucose concentrations with or without the addition of 20mM L-lactate. (C) Schematic of cytosolic LDH catalyzed oxidation of L-lactate and L-alpha-hydroxybutyrate to their respective alpha-ketoacids. (D) Glucose consumption of HepG2 cells cultured in medium with 5mM glucose with the addition of NaCl (control), L-lactate, or L-AHB at 20mM each. (E) Glycolytic proton efflux rate (GlycoPER) as measured using the Seahorse Bioanalyzer of HepG2 cells cultured in medium with 5mM glucose with the addition of NaCl (control), L-lactate, or L-AHB at 20mM each. (F) Media glucose consumption of HepG2 cells containing a doxycycline (Dox) inducible *Lb*NOX vector following the indicated treatments with or without the addition of 20mM L-lactate in medium containing 5mM glucose. (G) Oxygen consumption rate (OCR) of HepG2 cells measured using Seahorse Bioanalyzer. Metabolite arrow indicates injection of either 20mM NaCl (control) or 20mM L-lactate. Oligo, oligomycin; Rot/AA, rotenone/antimycin A. (H) Glucose consumption of HepG2 cells cultured in 2mM glucose following the indicated treatment with or without the addition of 10mM L-lactate. Untreated (-) or control conditions in this and subsequent figures indicate equimolar NaCl. All error bars represent mean +/± SD with a minimum n of 3. Statistical analysis in (B), (F), and (H) was performed using two-sided Student’s t-tests, in (D) and (E) was performed using one way ANOVA. **** p < 0.0001, * p < 0.05, and ns nonsignificant. See also Figure S1.

To investigate the possibility that suppression of glycolysis is due to reduction of cytosolic NAD^+^/NADH ratio through LDH-mediated lactate-to-pyruvate conversion, we asked whether L-alpha-hydroxybutyrate (AHB), an LDH substrate and cytosolic redox equivalent of L-lactate, had similar effects^13^ (Figure 1C). L-AHB treatment did not result in glycolytic suppression as measured by cellular glucose consumption and glycolytic proton efflux rate (glycoPER) (Figures 1D and 1E). Conversely, increasing the cytosolic NAD^+^/NADH ratio by expressing *Lb*NOX, a *Lactobacillus brevis* NADH oxidase that directly converts cytosolic NADH to NAD^+^ did not suppress lactate’s effects on glucose metabolism^14^ (Figures 1F, S1C, and S1D). These results suggest modulating cytosolic redox alone was insufficient to explain lactate’s suppression of glycolysis.

### Lactate stimulates mitochondrial ATP production

Next, we assessed the effects of lactate on mitochondrial respiration. When added to the medium, lactate rapidly increased the cellular oxygen consumption rate (OCR) (Figure 1G). When the ATP synthase inhibitor oligomycin was added, the OCR of both control (equimolar NaCl) and L-lactate treated cells declined to the same basal level. The addition of a mitochondrial uncoupling agent FCCP restored OCR in both conditions. Treatment with the respective ETC complex I and III inhibitors, rotenone and antimycin A, abolished the effect of lactate and resulted in a similar baseline OCR in both conditions. These results indicate lactate stimulates mitochondrial ATP-coupled ETC activity and prompted us to examine the role of increasing mitochondrial ATP production on lactate’s ability to suppress glycolysis. The addition of as little as 5nM of oligomycin to the culture medium was sufficient to reverse lactate-mediated glycolysis suppression (Figure 1H). The ability of oligomycin to reverse lactate-mediated glycolysis suppression was independent of any changes to mitochondrial NAD^+^/NADH ratio as it is unaffected by the further addition of FCCP, which enables maximal mitochondrial respiration and NAD^+^/NADH ratio. These results indicate that lactate suppresses glycolysis through increased mitochondrial ATP production rather than by modulating NAD^+^/NADH ratio alone.

### Lactate can mediate the Pasteur effect in mammalian cells independent of LDH

Known as the Pasteur effect, the ability of oxidative phosphorylation to suppress glucose fermentation was first described over 150 years ago during studies of yeast^15^. In mammals, the Pasteur effect has been tied to oxygen availability. But the mechanism by which oxidative phosphorylation suppresses glycolysis remains poorly understood^16^. It was unexpected that the glycolytic end product, lactate, might contribute to the ability of oxidative phosphorylation to suppress glycolysis. We therefore set out to test whether lactate has metabolic regulatory functions beyond its known role as a metabolic substrate.

Studies of metabolic regulatory functions of lactate can be confounded by its role in cytosolic redox homeostasis and as a carbon source. These known metabolic roles of lactate are dependent on LDH catalyzed reactions. However, lactate can exist in two enantiomeric forms and the metabolism through mammalian LDHA/B is stereoselective for L-lactate or the (*S*)-enantiomers of alpha-keto acids^17^ (Figure 2A). We therefore asked whether D-lactate, which is not an LDHA/B substrate, had similar effects. While a mammalian LDHD enzyme that’s expected to convert D-lactate to pyruvate has been cloned, its expression is tissue-selective, being expressed primarily in primary hepatocytes and is not present in most tissues^18, 19^. Consistently, we found that LDHD transcription expression as assessed by RNA-sequencing was extremely low to undetectable in our cell lines (Figures S2A-D). To measure LDHD enzymatic activity in cells, we performed tracing experiments using uniformly ^13^C labeled [U-^13^C] D-lactate. No pyruvate or citrate labeling was observed in multiple cell lines and non-transformed cells (Figures 2A, 2B, and S2E-H). Thus, LDHD activity does not contribute to NADH production or the generation of TCA cycle intermediates in the cells used in our study.

**Figure 2.**
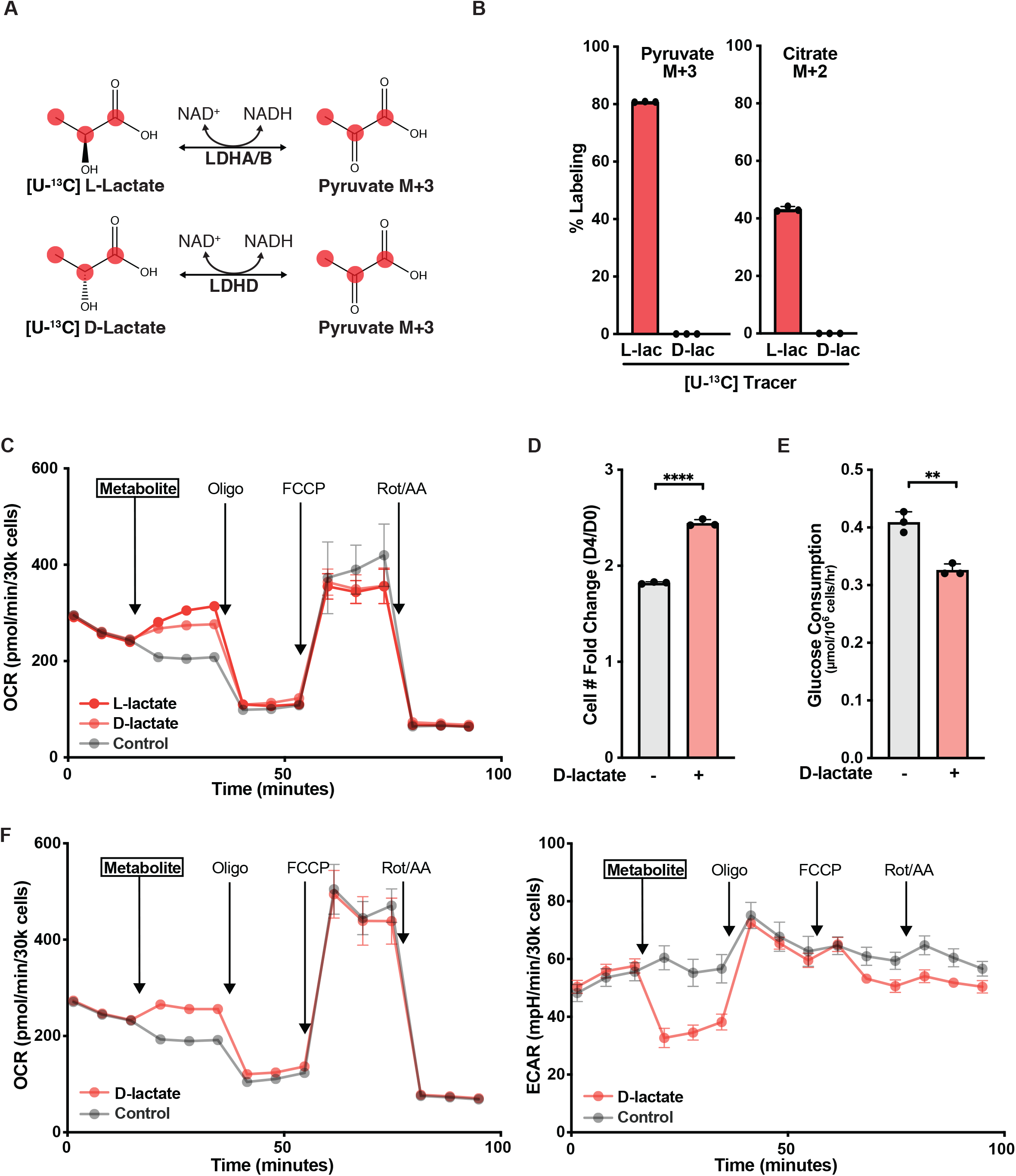
Both stereoisomers of lactate increase oxidative phosphorylation and lactate can suppress glycolysis independent of its metabolism. (A) Schematic of [U-^13^C] L-or D-lactate tracing to pyruvate through expected activities of LDHA/B/D. (B) Percent labeling of pyruvate M+3 or citrate M+2 in HepG2 cells following incubation with 10mM [U-^13^C] L- or D-lactate for 8 hours. (C) OCR of HepG2 cells measured using Seahorse Bioanalyzer. Metabolite arrow indicates injection of NaCl (control), L- or D-lactate at 20mM each. (D) Proliferation of HepG2 cells cultured in 1mM glucose with or without the addition of 20mM D-lactate measured by cell number fold change relative to day 0. (E) Glucose consumption of HepG2 cells cultured in medium with 5mM glucose with or without the addition of 20mM D-lactate. (F) OCR and ECAR of HepG2 cells measured using Seahorse Bioanalyzer. Metabolite arrow indicates injection of NaCl (control) or D-lactate at 20mM each. All error bars represent mean +/± SD with a minimum n of 3. Statistical analysis in (D) and (E) was performed using two-sided Student’s t-tests. ** p < 0.01, **** p < 0.0001. See also Figure S2.

Strikingly, despite not being metabolized to pyruvate or citrate, D-lactate addition to the medium rapidly increased mitochondrial OCR and coupled ATP production (Figure 2C). Consistently, the addition of D-lactate increased the survival and proliferation of cells in low glucose and suppressed cellular glucose consumption (Figures 2D and 2E). Collectively, our results demonstrate that both stereoisomers of lactate can increase oxidative phosphorylation and suppress glycolysis, and these effects are independent of lactate conversion to pyruvate or entry into the TCA cycle.

We next tested the ability of D-lactate to reverse aerobic glycolysis. As shown in Figures 2B and 2C, D-lactate stimulates oxygen consumption and mitochondrial ATP production without being converted to pyruvate or citrate. When the extracellular acidification rate (ECAR) was simultaneously measured (Figure 2F), we found D-lactate simulation of oxygen consumption correlated with a reduction of extracellular acidification. More importantly, when mitochondrial ATP production was inhibited with oligomycin, there was a coincident recovery of glycolysis to the level of control cells as measured by a recovery in the extracellular acidification rate. These results indicate that D-lactate can suppress glucose fermentation through stimulating oxidative phosphorylation and that aerobic glycolysis (Warburg effect) is reversible.

### Lactate stimulation of mitochondrial respiration increases pyruvate entry into the TCA cycle

The ability of lactate to increase oxidative phosphorylation prompted us to examine its effects on carbon entry into the TCA cycle, which is tightly coupled to cellular oxidative phosphorylation and ETC activity^20^. Pyruvate is a major carbon that enters the TCA cycle through the pyruvate dehydrogenase (PDH) complex. Both glucose and lactate can produce pyruvate, though their relative contributions are unclear. We used isotope tracing with [U-^13^C] glucose and [3-^13^C] L-lactate to simultaneously determine the fate of lactate and glucose carbon across varying lactate concentrations (Figure 3A). This analysis revealed lactate to be the predominant cellular source of pyruvate even when glucose is in vast excess, and lactate’s contribution to pyruvate increased in a dose-dependent fashion (Figures 3B, 3C, S3A, and S3B). Furthermore, pyruvate flux into the TCA cycle increased in a dose-dependent fashion as the lactate concentration increased. These results prompted us to examine lactate’s effect on activity of the PDH complex, the rate limiting step for pyruvate entry into the TCA cycle.

**Figure 3.**
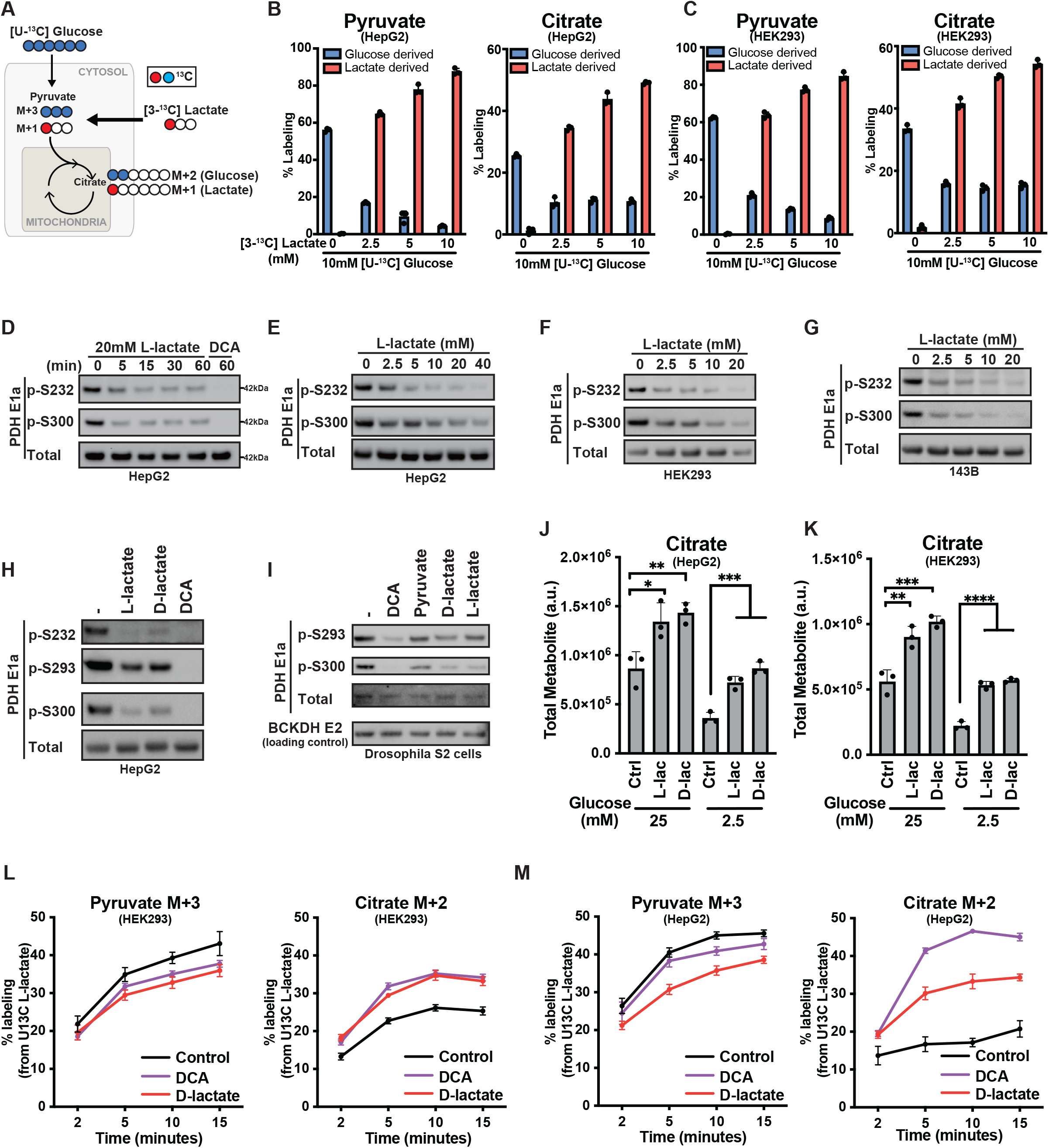
Lactate stimulation of mitochondrial respiration increases use of pyruvate as a TCA substrate. (A) Schematic of [U-^13^C] glucose and [3-^13^C] lactate tracing into the TCA. (B and C) Percent labeling of pyruvate and citrate from 10mM [U-^13^C] glucose with indicated concentration of [3-^13^C] lactate in HepG2 and HEK293 cells. (D) HepG2 cells were treated for the indicated time with 20mM L-lactate or 5mM DCA followed by immunoblotting analysis with the indicated antibodies. (E) HepG2, (F) HEK293, and (G) 143B cells were treated with indicated L-lactate concentrations followed by immunoblotting analysis with the indicated antibodies. (H) HepG2 cells were treated as indicated (20mM NaCl, L- or D-lactate, or 5mM DCA) followed by immunoblotting analysis. (I) Drosophila S2 Cells were treated as indicated (20mM NaCl, 5mM DCA, 2mM pyruvate, and 20mM D- or L-lactate) followed by immunoblotting analysis. (J) HepG2 and (K) HEK293 cells cultured in medium containing 25mM or 2.5mM glucose were incubated with 20mM NaCl, L- or D-lactate for 1 hour before total citrate abundance was analyzed using GC-MS. (L) HepG2 and (M) HEK293 cells cultured in 25mM glucose and 2.5mM [U-^13^C] lactate were treated with 20mM NaCl, 5mM DCA, or 20mM D-lactate and the % ^13^C labeling of pyruvate and citrate from [U-^13^C] lactate was analyzed over the time course using GC-MS. All error bars represent mean +/± SD with a minimum n of 3. Statistical analysis in (J) and (K) was performed using one way ANOVA. *p < 0.05, **p < 0.01, ***p < 0.001, **** p < 0.0001. See also Figure S3.

PDH activity is regulated by the levels of inhibitory phosphorylation on the E1α subunit at three serine residues: S232, S293, and S300. PDH E1α phosphorylation is balanced by the activities of pyruvate dehydrogenase kinases (PDKs) and phosphatases (PDPs) and is modulated by the mitochondrial ETC activity and NAD^+^/NADH ratio^20^. Cells grown in complete medium had significant PDH E1α phosphorylation that could be reduced by the PDK inhibitor dichloroacetate (DCA), which directly activates the PDH complex^21^ (Figures 3D). Furthermore, increasing ETC activity and mitochondrial NAD^+^/NADH using the uncoupler FCCP led to expected PDH activation, as indicated by decreased inhibitory E1α phosphorylation (Figures S3C to S3E).

Suppressing ETC and decreasing mitochondrial NAD^+^/NADH with a complex I or III inhibitor, rotenone or antimycin A, respectively, led to PDH inactivation with an increase in E1α phosphorylation (Figures S3C to S3E). The addition of L-lactate to the medium resulted in a dose-and time-dependent activation of PDH, as shown by a decrease in E1α inhibitory phosphorylation across multiple serine residues and across multiple cell lines (Figures 3D to 3G and S3F to S3H).

Consistent with an LDH-independent mechanism, D-lactate was comparable to L-lactate in activating PDH and reducing its phosphorylation in cells (Figure 3H). This ability was evolutionarily conserved. Drosophila S2 cells also exhibited suppression of PDH Elα phosphorylation when treated with D- or L-lactate (Figure 3I).

To confirm that lactate-induced activation of PDH increased carbon entry into the TCA cycle, we first measured the absolute amount of both pyruvate and citrate after cells were treated with either D- or L-lactate. Despite not being metabolized to pyruvate or citrate, D-lactate significantly increased total levels of citrate in either high (25mM) or low (2.5mM) glucose medium similarly as L-lactate, while total cellular pyruvate levels remained minimally affected (Figures 3J, 3K, S3I and S3J). As our earlier tracing results suggested lactate to be the predominant cellular pyruvate source, an increase in pyruvate oxidization through PDH activation is predicated to also increase lactate oxidation. We therefore assessed whether D-lactate, which contains lactate’s metabolic regulatory but not catabolic function, can increase the entry of L-lactate carbons into the TCA cycle. In cells in complete medium supplemented with 2.5 mM [U-^13^C] L-lactate, the addition of D-lactate significantly increased the rate of citrate M+2 labeling from [U-^13^C] L-lactate even though L-lactate labeling of pyruvate (M+3) was slightly reduced (Figures 3L and 3M). These results demonstrate that independent of its metabolism, lactate stimulation of mitochondrial respiration leads to an increase of pyruvate entry and oxidation into the mitochondrial TCA cycle.

### Lactate addition activates the PDH complex in isolated mitochondria

As our results indicate lactate can activate PDH and increase TCA flux independent of its cytosolic functions, we sought to dissect lactate’s function on isolated mitochondria in a cell-free assay using PDH activation as a readout. Mitochondria purified using established cell fractionation procedures were incubated in an isotonic buffer with respiratory substrates (Figure 4A)^22, 23^. As expected for coupled mitochondria, ADP stimulated ETC activity and reduced PDH phosphorylation while ATP addition reduced ETC activity and increased inhibitory phosphorylation of PDH (Figure 4B). As a positive control, the kinase inhibitor DCA led to the dephosphorylation of Elα subunits of both the PDH and the branch chain ketoacid dehydrogenase (BCKDH) complex, a closely related mitochondrial α-ketoacid dehydrogenase family member that is known to be activated also by DCA^24^. In contrast, L-lactate addition led to selective dephosphorylation of PDH Elα and not BCKDH Elα. The ability of L-lactate to reduce PDH phosphorylation in purified mitochondria was greater than that of pyruvate.

**Figure 4.**
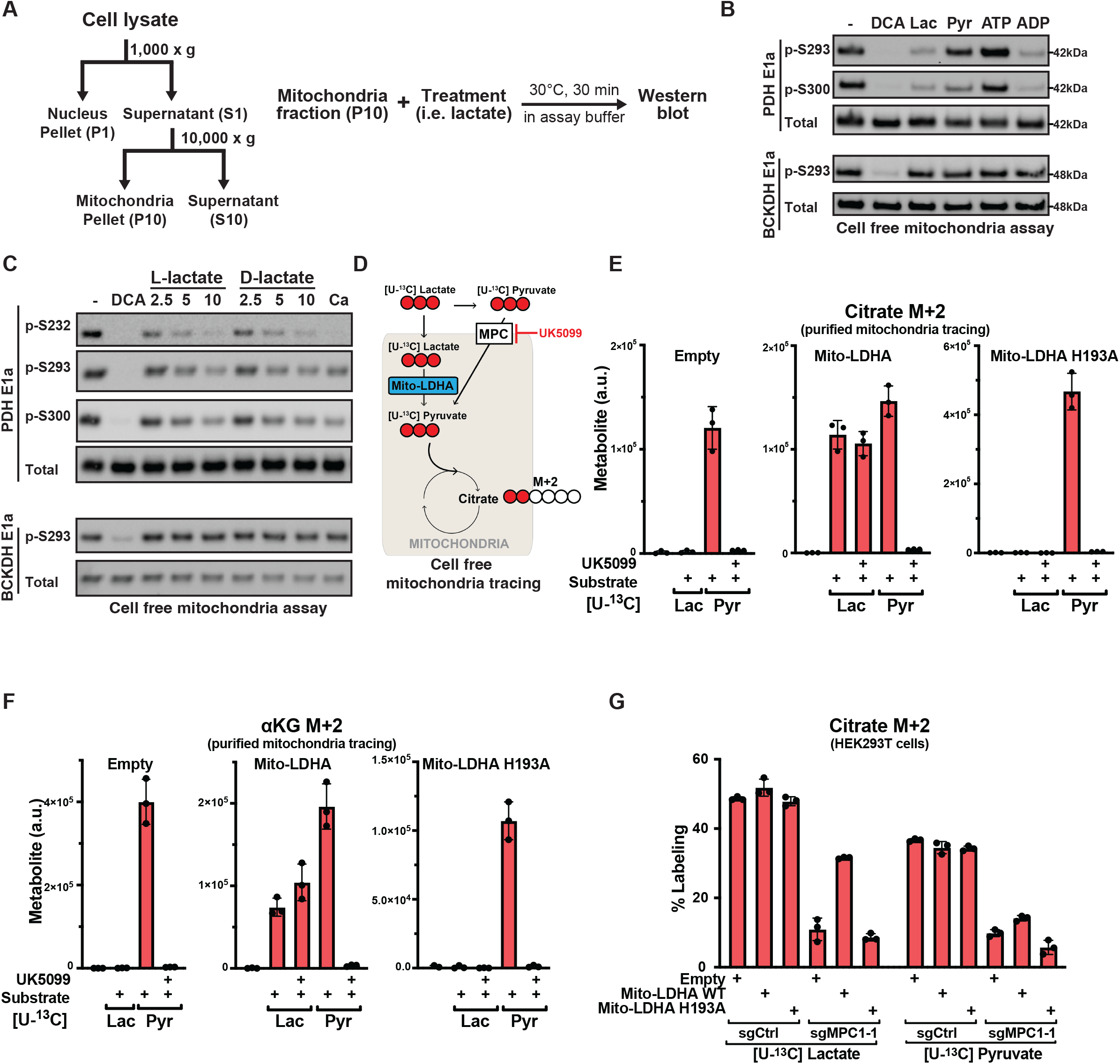
Lactate activates the PDH complex in isolated mitochondria in a dose dependent manner and directly enters the mitochondria matrix independent of mitochondrial pyruvate carrier. (A) Schematic of subcellular fractionation and cell free assay using purified mitochondria. (B) Mitochondria purified from 293T cells were incubated in assay buffer with the indicated treatments (5mM NaCl, DCA, L-lactate, pyruvate, 4mM ATP or ADP) for 30 mins at 30°C, followed by immunoblotting with the indicated antibodies. (C) Purified mitochondria were incubated with indicated concentration of L- or D-lactate (mM), 5mM DCA, or 5mM calcium (Ca, phosphatase activator) followed by immunoblotting analysis. (D) Schematic of [U-^13^C] pyruvate or [U-^13^C] lactate tracing in purified mitochondria. (E and F) Mitochondria purified from 293T cells expressing empty vector or mitochondrial-matrix targeted (Mito-) wildtype or mutant (H193A) LDHA were incubated with or without substrate (2mM [U-^13^C] lactate or 2mM [U-^13^C] pyruvate) in the presence or absence of the MPC blocker UK5099 (10µM), followed by GC-MS analysis of total citrate and α-ketoglutarate M+2 abundance in the reaction mixture. (G) 293T cells expressing mitochondrial-matrix targeted (Mito-) LDHA WT or H193A with or without MPC1 deletion were cultured in media containing 10mM [U^13^C] L-lactate or 2mM [U^13^C] pyruvate followed by GC-MS analysis of citrate M+2 labeling. All error bars represent mean ± SD with a minimum n of 3. See also Figure S4.

When the stereospecificity of lactate activation of PDH was examined in purified mitochondria, L-lactate and D-lactate were found to be equivalent in their ability to suppress PDH Elα-phosphorylation, without affecting BCKDH E1α (Figure 4C). Our cell-free mitochondria assay eliminates variable transport across the plasma membrane as L-lactate has a lower Km for the monocarboxylate transporters (MCTs) than D-lactate^25^. While both D- and L-lactate can support PDH activation, both the D- and L- enantiomers of alanine, which contains an amino group in place of lactate’s hydroxyl group, were unable to activate PDH (Figure S4A), indicating some specificity to lactate’s ability to activate PDH.

### Lactate directly enters the mitochondrial matrix independent of the mitochondrial pyruvate carrier (MPC)

The ability of lactate to stimulate mitochondrial activity in isolated mitochondria prompted an examination of lactate metabolism within and transport into the mitochondrial matrix^26^. We first performed carbon tracing in wildtype mitochondria, where [U-^13^C] pyruvate led to expected TCA substrate labeling that was abolished when UK5099, the MPC inhibitor was added (Figures 4D, S4B, and 4E and 4F, left panels). The addition of uniformly labeled [U-^13^C] L-lactate to purified wildtype mitochondria did not result in citrate or α-ketoglutarate (αKG) labeling, indicating lactate cannot be directly metabolized in purified mitochondria. These results indicate that like D-lactate, the ability of L-lactate to activate the PDH complex in purified mitochondria is also independent of its metabolism.

Next, we performed isotope tracing in isolated mitochondria expressing active or inactive forms of mitochondrial matrix-targeted LDHA (mito-LDHA) (Figures 4D and S4B). In contrast to wildtype mitochondria, when [U-^13^C] L-lactate was added to mitochondria expressing mito-LDHA, significant citrate and αKG labeling was observed (Figures 4E and 4F, middle panels). Notably, unlike pyruvate, lactate labeling of citrate and αKG was not reduced by UK5099 and occurred only in mitochondria expressing mito-LDHA but not the H193A catalytic LDHA mutant (Figures 4E and F, right panels). Similar results were obtained in intact cells (Figure 4G), where deletion of MPC1 in control cells suppressed pyruvate entry into the TCA cycle.

However, in cells also expressing a functional mito-LDHA, L-lactate retained the ability to label TCA cycle citrate and αKG even in the absence of MPC (Figures 4G and S4C). Collectively, these results indicate that lactate can enter the mitochondrial matrix, consistent with the recent demonstration of the existence of lactate in the mitochondrial matrix using imaging probes^27^. Moreover, our results indicate that lactate’s mitochondrial entry is direct and not due to secondary metabolism in the matrix and that lactate entry is independent of the MPC.

Consistent with the observed effect of lactate on mitochondria being MPC-independent, deletion of the MPC1 did not affect the ability of either L-lactate or D-lactate to induce mitochondrial oxidative phosphorylation, while the effects of pyruvate addition were significantly inhibited by MPC1 deletion (Figures S4C and S4D). Similarly, lactate but not pyruvate retained its ability to activate PDH following UK5099 treatment (Figure S4E). These results indicate that lactate stimulates the activity of purified mitochondria without being metabolized to pyruvate and directly enters the mitochondrial matrix independent of MPC.

### Lactate activates the electron transport chain

Next, we sought to determine the sequence of lactate’s action on the mitochondrial ETC and PDH complex. Lactate could directly activate PDH to increase pyruvate influx with subsequent increase in ETC activity as a secondary effect. We reasoned that if lactate had a direct allosteric effect on the PDH complex, then that effect should be independent of mitochondrial membrane integrity. However, when purified mitochondria were permeabilized by repeated free/thaw cycles or with mild non-ionic detergents, the ability of both D-lactate and L-lactate to suppress PDH Elα phosphorylation was abolished (Figures 5A and 5B). In contrast, the effects on PDH Elα phosphorylation of PDK-inhibitor DCA or the PDP-activator calcium were retained in permeabilized mitochondria. These results indicate that lactate is unlikely to directly activate the PDH complex.

**Figure 5.**
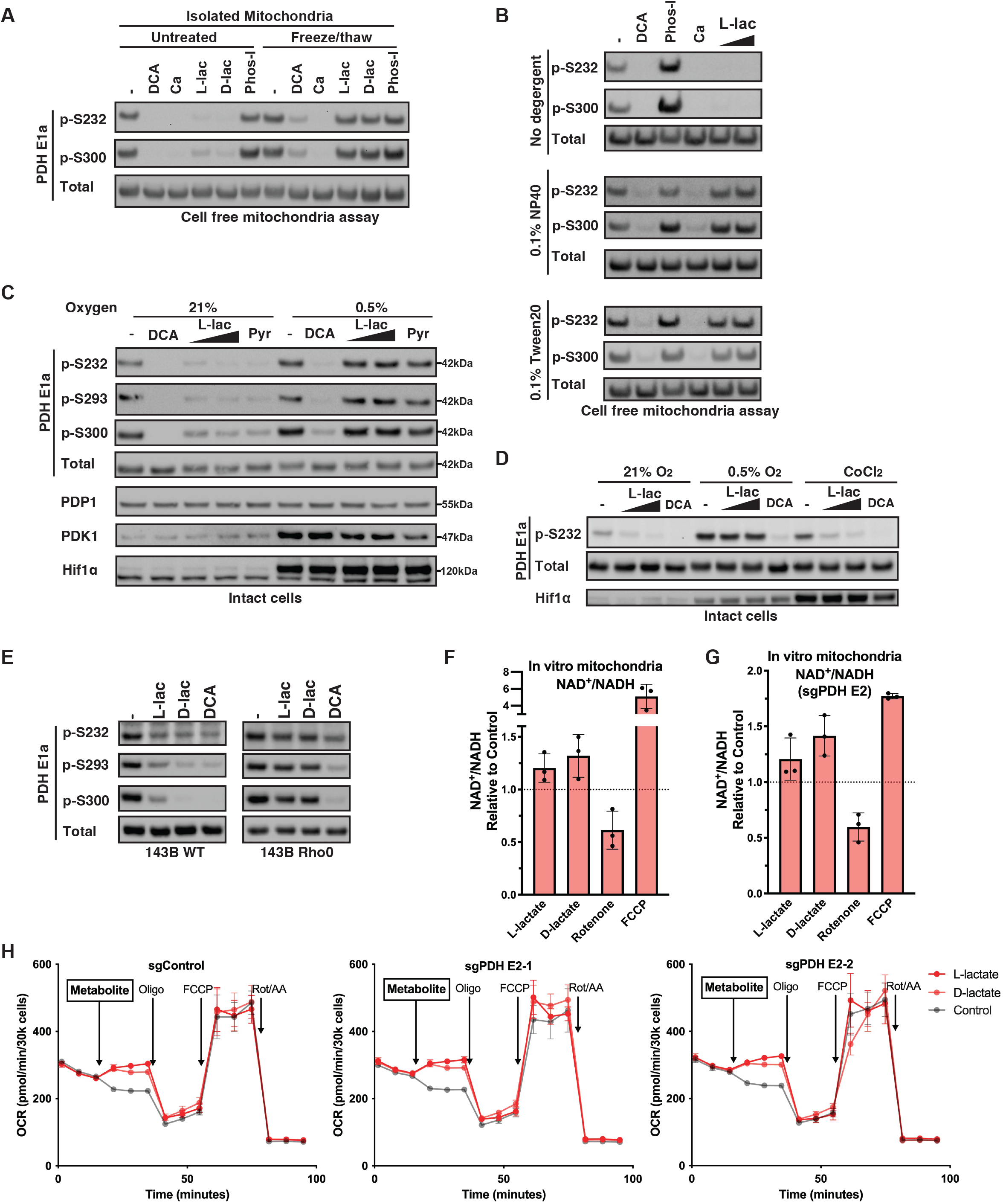
Lactate signals through the electron transport chain independent of mitochondrial pyruvate entry. (A) Purified mitochondria were subjected to rapid freeze and thaw or left on ice (untreated), followed by incubating at 30°C for 30 mins in assay buffer with the indicated treatments at 5mM followed by immunoblotting analysis. Ca, calcium; Phos-I, phosphatase inhibitor at 1x. (B) Purified mitochondria were resuspended in assay buffer with or without the indicated detergent, followed by cell-free mitochondria assay as described in (A) with the indicated treatments. Phos-I was at 1x, L-lactate was used at 5mM and 15mM, and all others at 5mM. (C) Following overnight culture at the indicated oxygen concentration, HepG2 cells were treated with 5mM DCA, 15mM or 30mM L-lactate, or 2mM pyruvate for 30 mins at the indicated oxygen concentration, followed by immunoblotting. (D) Following overnight culture at the indicated oxygen concentration or in 100µM CoCl2, HepG2 cells were treated with 15mM or 30mM lactate or 5mM DCA for 30 mins followed by immunoblotting. (E) 143B Rho0 or matched WT control cells were treated with 20mM L-/D-lactate or 5mM DCA for 30 mins followed by immunoblotting. (F and G) Purified mitochondria from 293T or 293T sgPDH E2-1 cells were incubated in assay buffer containing 10mM NaCl (control), 10mM L-/D-lactate, 1µM Rotenone, or 1µM FCCP for 30 mins at 30°C, followed by measuring the total NAD^+^ and NADH levels in the reaction using a modified enzyme cycling assay. (H) OCR of HepG2 cells containing the indicated sgRNA measured using Seahorse Bioanalyzer. Metabolite arrow indicates injection of either 20mM NaCl (control), L- or D-lactate. All untreated or control conditions indicate NaCl equimolar to that of the highest lactate concentration. All error bars represent mean ± SD with a minimum n of 3. See also Figure S5.

We next considered the possibility that lactate stimulated ETC activity first and decreased PDH phosphorylation as a downstream effect. As the predominant electron acceptor in cells, oxygen plays an essential role in regulating ETC activity and mitochondrial redox^28^. Culturing cells overnight under hypoxic conditions (0.5% oxygen) abolished the ability of lactate addition to reduce PDH Elα phosphorylation (Figures 5C and S5A). This effect was independent of Hif1α stabilization as CoCl2 treatment stabilized Hif1α but did not affect lactate-induced PDH Elα dephosphorylation (Figures 5D and S5B). These results indicate that the ability of lactate to reduce PDH Elα phosphorylation depended on oxygen availability rather than being suppressed by Hif1α.

Next, to directly test the role of the ETC, we turned to Rho0 cells, which are deficient in mitochondrial DNA and therefore lack a functional ETC^29, 30^. Compared to the wildtype 143B cells, DCA retained its ability to dephosphorylate PDH Elα in ETC-deficient Rho0 cells. However, L- and D-lactate’s ability to induce PDH Elα dephosphorylation was lost in the Rho0 cells (Figure 5E), indicating the ETC is essential in lactate-induced PDH activation.

The mitochondrial ETC converts NADH to NAD^+^, and an increase in mitochondrial NAD^+^/NADH ratio is known to activate the PDH complex^14^. Incubation of isolated mitochondria with lactate resulted in an increase in mitochondrial NAD^+^/NADH ratio (Figure 5F). Consistent with an ability to activate the ETC independent of mitochondrial pyruvate entry, lactate increased NAD^+^/NADH ratio even in mitochondria deficient in pyruvate dehydrogenase complex (PDH E2 deletion) (Figures 5G and S5C). Consistently, lactate retained the ability to increase mitochondrial OCR in cells with PDH E1α or E2 deficiency, indicating the increase in ETC activity was independent of carbon entry through the PDH complex and that PDH activation is a secondary effect of lactate activation of the ETC (Figures 5H and S5D to S5H).

### D-lactate enables proliferation of cells with compromised respiration

The above results indicate that L- and D-lactate exert similar effects on mitochondrial oxidative phosphorylation, PDH activation, and TCA cycle flux. However, they differ in their cytosolic effects. While D-lactate is not metabolized to pyruvate in the cell lines studied, L-lactate through

LDH mediated metabolism generates both pyruvate and NADH in the cytosol. The excessive cytosolic NADH produced by L-lactate metabolism can lead to reductive stress, elevated reactive oxygen species, and impaired synthesis of serine and nucleotide^31–34^. For example, T cell treatment with L-lactate has been reported to suppress glycolytic production of serine by reducing cytosolic NAD^+^ levels^34^. Consistent with this, we found L-lactate suppressed the growth of T-cells in serine-deficient media. In contrast, D-lactate supported and even enhanced cell growth in serine-deficient medium (Figure S6A). Therefore, to further examine lactate’s non-LDH dependent aspect as a mitochondrial messenger, we focused on further characterizing lactate’s effect on mitochondrial oxidative phosphorylation and electron transport using D-lactate.

Given its ability to stimulate oxidative phosphorylation, we first asked whether D-lactate can enhance the proliferation of cells dependent on oxidative phosphorylation for ATP production. A long sought after goal in the field of cellular bioengineering has been to grow cells purely on oxidizable substrates, such as galactose. Cells cultured in galactose are fully dependent on the mitochondria for ATP production^35^. Most mammalian cells cultured in medium containing galactose as their only source of monosaccharide lose their ability to proliferate and/or die^36, 37^. However, we found that addition of D-lactate rescued cell survival and promoted cell proliferation in both transformed and non-transformed cells (Figures 6A and 6D).

**Figure 6.**
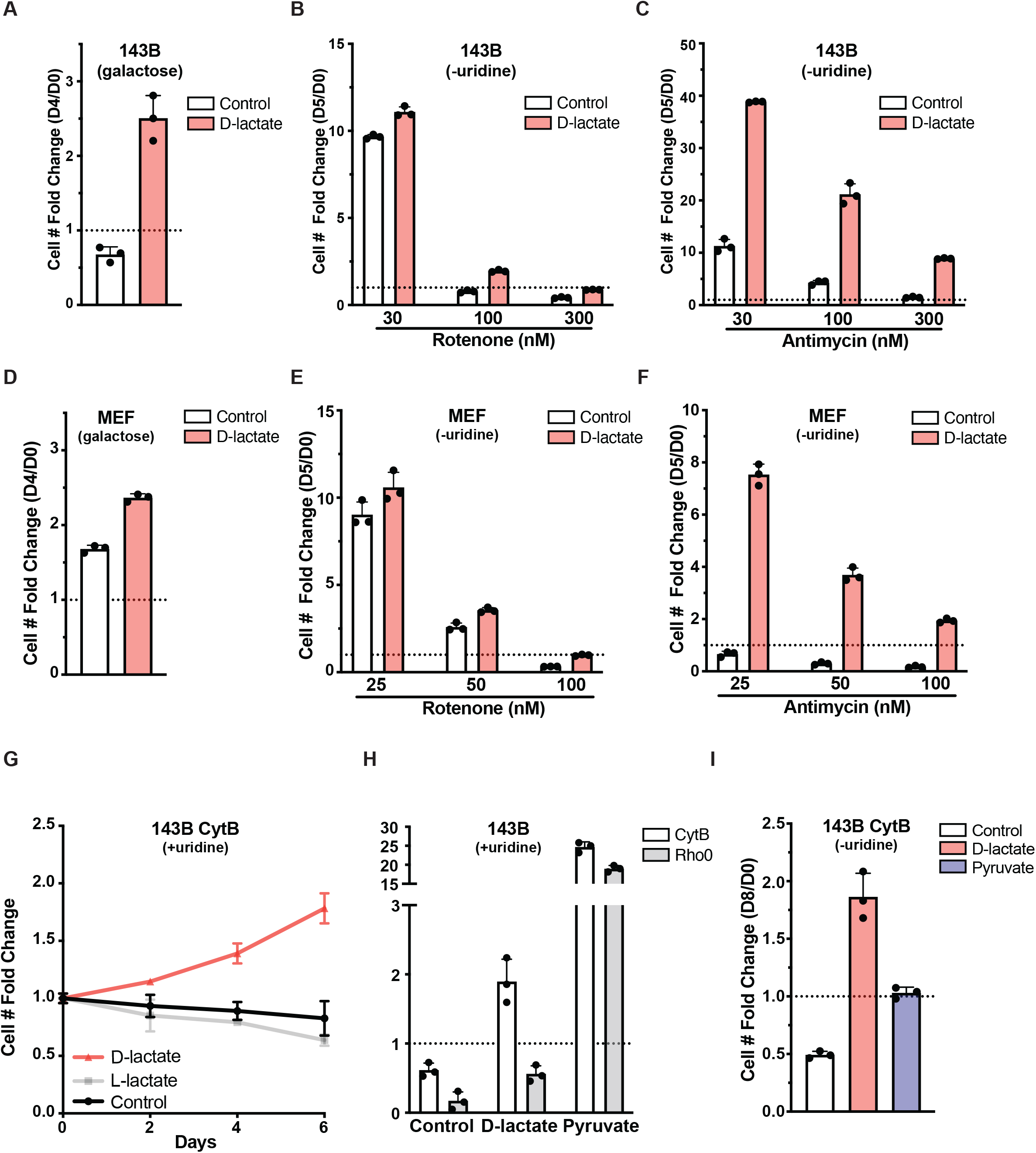
D-lactate enhances respiration dependent cell proliferation. (A to F) 143B and MEF cells were cultured in glucose-deficient DMEM supplemented with galactose (A and D) or complete DMEM containing the indicated rotenone (B and E) or antimycin A (C and F) concentrations, along with the addition of 20mM NaCl or D-lactate. Proliferation was measured by cell number fold change relative to day 0. (G) Proliferation of 143B cytochrome-B cybrids (143B CytB) cultured in complete DMEM with uridine and supplemented with 20mM NaCl, L-lactate, or D-lactate measured as cell number fold change relative to day 0. (H) Proliferation of 143B CytB or 143B Rho0 cells cultured in DMEM with uridine and the indicated treatments was measured as cell number fold change relative to day 0. (I) Proliferation of 143B CytB cells cultured in complete DMEM with the indicated treatment conditions in the absence of uridine. All error bars represent mean +/± SD with a minimum n of 3. See also Figure S6.

We next examined the effect of D-lactate on cells with impaired ETC function. Respiration defective cells have impaired proliferation and require supplemental electron acceptors to enable biosynthesis and ATP production, most often in the form of pyruvate, to survive and proliferate^13, 14, 38, 39^. We examined D-lactate’s effect on the proliferation of cells cultured in increasing doses of rotenone (Complex I inhibitor) or antimycin A (Complex III inhibitor). While D-lactate provided a modest growth advantage in rotenone, it significantly enhanced the ability of cells to proliferate when treated with antimycin A (Figures 6B, 6C, 6E, and 6F).

The ability of D-lactate to stimulate coupled oxidative phosphorylation and rescue the effects of ETC inhibitors suggests it may enhance electron transfer efficiency. To examine this possibility, the levels of mitochondrial reactive oxygen species (ROS) in rotenone- or antimycin-treated cells was assessed. Mitochondrial superoxide generation was reduced by treatment with D-lactate to levels comparable to that provided by pyruvate (Figure S6B), suggesting D-lactate’s ability to selectively stimulate the ETC can prevent the build-up of excessive NADH.

To further investigate the selective effects of lactate on ETC activity, we studied the effects of D-lactate on 143B CytB cells, which are cybrid cells containing a patient-derived 4-bp deletion in the cytochrome B gene that is a component of Complex III, and 143B Rho0 cells that are completely devoid of mitochondrial DNA^30, 40^. Unlike Rho0 cells that are completely deficient in ETC activity, 143B CytB mutant cells have residual electron transport activity^40–42^. As opposed to the parental 143B cells, both 143B CytB cells and 143B Rho0 cells normally require both uridine and pyruvate supplementation to maintain viability and growth^38, 40^. Pyruvate supports the metabolic adaptation of respiration defective cells by increasing cytosolic NAD^+^ through the LDH reaction. As expected, L-lactate, which has the opposite cytosolic redox effect to pyruvate, was unable to support 143B CytB proliferation (Figure 6G). However, D-lactate enabled the continued proliferation of 143B CytB cells in the absence of pyruvate, consistent with its role in selectively increasing ETC activity without being metabolized through cytosolic LDH and affecting cytosolic redox (Figure 6G). While D-lactate can support the proliferation of 143B CytB cells, it was unable to maintain the survival or proliferation of 143B Rho0 cells, suggesting that D-lactate’s ability to promote survival and growth required some ETC activity (Figure 6H).

As 143B CytB cells have elevated ROS due to incomplete electron transfer at complex III, we asked whether D-lactate can decrease mitochondrial ROS in these cells. D-lactate, despite not being metabolized in 143B cells, reduced mitochondrial ROS to levels comparable to pyruvate (Figure S6C). Finally, the effect of D-lactate on the uridine requirement of 143B CytB cells was distinct from that of pyruvate. Pyruvate supplementation kept 143B CytB cells alive but the cells could not proliferate unless the medium contained supplemental uridine, the synthesis of which is dependent on functional electron transfer from DHODH through the Q-cycle to complex III.

In contrast, D-lactate enabled modest proliferation of 143B CytB cells in the absence of uridine (Figure 6I and S6D). These data suggest that D-lactate stimulation of ETC activity not only enhances the transfer of electrons donated by the TCA cycle within matrix but also electrons transferred by reaction found outside of the mitochondrial matrix such as those produced by DHODH activity.

### D-lactate enhances primary T-cell proliferation and effector function

The ability of lactate to enhance cellular respiration prompted us to examine its role in improving T-cell proliferation and function, which can be limited by their oxidative phosphorylation capacity. T-cells are being routinely expanded in culture for use in cellular immunotherapy and gene therapy^43^. One persistent challenge that limits the efficacy of such T cells is termed T cell exhaustion, which is associated with impaired oxidative phosphorylation and reduced ATP/GTP levels^44, 45^. L-lactate has been shown to impair T-cell proliferation due to its metabolism through

LDH, which results in an increase in cytosolic NADH that impairs serine biosynthesis^34^. We therefore asked whether D-lactate, which bypasses cytosolic LDH metabolism and does not impair T-cell growth in serine deficient medium (Figure S6A), can improve T cell mitochondrial effector function and/or growth. Freshly isolated T cells were expanded with a combination of αCD3 and αCD28 supplemented with interleukin 2 in complete medium with or without D-lactate. The addition of D-lactate stimulated mitochondrial respiration and increased coupled ATP synthesis (Figure 7A). D-lactate also suppressed T-cell glucose consumption (Figure 7B). These results are consistent with our prior observation in immortalized cell lines and suggest that D-lactate can stimulate mitochondrial oxidative phosphorylation while reducing the high rate of glycolysis that leads to frequent media changes during T cell preparation for cellular immunotherapy.

**Figure 7.**
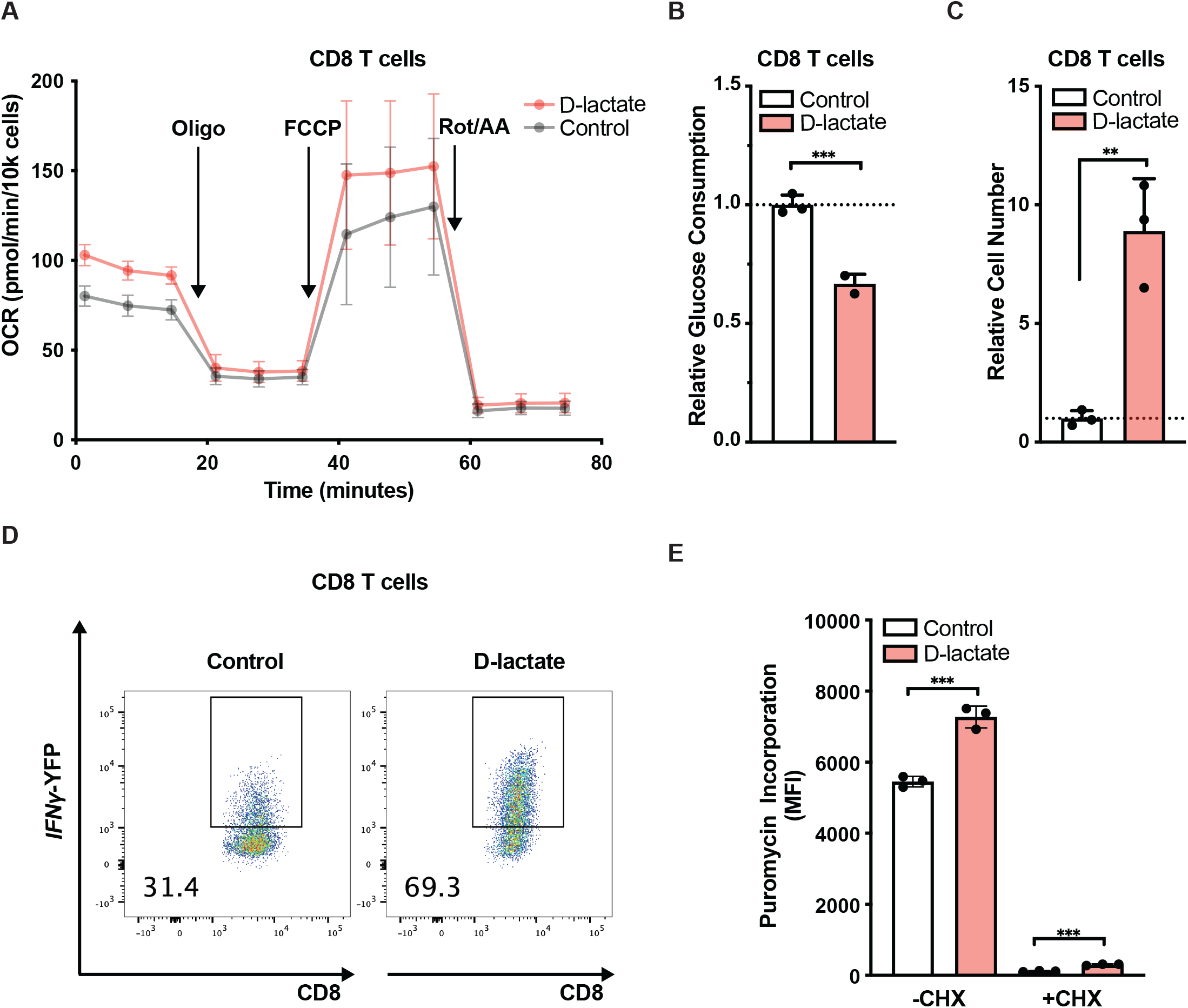
D-lactate enhances primary T-cell proliferation and effector function. (A) Mitochondrial OCR of primary murine CD8 T cells in standard RPMI with or without the addition of D-lactate as measured using Seahorse Bioanalyzer. (B) Glucose consumption of primary murine CD8 T cells cultured in standard RPMI media with or without the addition of 20mM D-lactate. (C) Relative cell number of primary murine CD8 T cells cultured in 1mM glucose with or without the addition of 20mM D-lactate. (D) IFN-γ production in CD8 T cells cultured in 1mM glucose with or without the addition of 20mM D-lactate. (E) Puromycin incorporation in CD8 T cells as determined by flow cytometry following NaCl (20mM) or Na-D-lactate (20mM) treatment. All error bars represent mean +/± SD with a minimum n of 3. Statistical analysis in (B), (C) and (E) was performed using two-sided Student’s t-tests. **p < 0.01, ***p < 0.001.

We next examined whether D-lactate can enhance T cell function under glucose limiting conditions that are often encountered in tumors. As reported previously, as glucose concentrations in the medium are reduced to 1mM, further T cell expansion and the maintenance of cytokine production become compromised^46, 47^. Under these conditions, D-lactate restored T cell proliferative expansion (Figure 7C). T cells increased eightfold in number and have a marked increase in the expression of interferon-γ, a marker of their effector function (Figure 7D). Consistent with its selective ability to stimulate the mitochondria, D-lactate was found to be superior to other metabolized monocarboxylates including L-lactate, pyruvate, acetoacetate, and β-hydroxybutyrate at inducing T-cell cytokine production (Figure S7). T-cell effector function is a highly ATP/GTP dependent process and we showed that D-lactate increases ATP-coupled ETC activity. Consistently, D-lactate treatment led to an increase in cellular protein synthesis rate, suggesting a mechanistic basis for the ability of D-lactate to stimulate T cell effector function is through ATP-dependent support of increased translation (Figure 7E).

## DISCUSSION

### Lactate acts as a mitochondrial messenger to simulate oxidative phosphorylation

Lactate is produced primarily through glycolysis. The glycolytic conversion of one glucose into two secreted molecules of lactate is redox neutral but yields 2 ATPs. As lactate is not appreciably excreted from the body, excess circulating lactate needs to either be oxidized to CO2 in the mitochondria or be used to build macromolecules. Mitochondrial oxidative phosphorylation produces ATP more efficiently than aerobic glycolysis. However, how cells regulate the relative contribution of aerobic glycolysis and oxidative phosphorylation is not understood. Furthermore, emerging studies have pointed to lactate, but not glucose, as the primary TCA cycle carbon source in mammals ^5–7, 48^. However, the mechanism by which lactate serves as the preferential carbon source for the mitochondria remains unclear.

Here we show that extracellular lactate acts in a dose-dependent fashion to increase oxidative phosphorylation and suppress glycolysis. Our results indicate that lactate acts as a messenger that activates the ETC to increase mitochondrial ATP production and pyruvate entry into the TCA cycle. As the LDH reaction is at equilibrium, the increase in mitochondrial pyruvate oxidation shifts the equilibrium between lactate and pyruvate to increase conversion of lactate to pyruvate, to support increased oxidative phosphorylation. The resulting increase in mitochondrial ATP production then suppresses glycolysis. Collectively, our results indicate that lactate stimulates its own use as a preferential bioenergetic substrate in the mitochondria as long as sufficient oxygen is available to support increase electron transport at the ETC.

While lactate is not essential for the ability of the mitochondria to perform oxidative phosphorylation, the results suggest lactate increases ETC flux and ATP production in a dose-dependent fashion. Lactate levels are known to fluctuate under physiological and pathological conditions^49–51^. Our results suggest extracellular lactate levels can serve to regulate and set basal oxidative phosphorylation activity under oxygen sufficiency. Lactate regulation of ETC activity and ATP-coupled oxygen consumption is distinct from other known stimulations of oxidative phosphorylation. Lactate stimulates ETC activity and ATP production while simultaneously reducing the mitochondrial NADH/NAD^+^ ratio and suppressing mitochondrial ROS generation. This is in contrast to other known mechanisms to increase ETC activity involving increasing flux into the TCA, which leads to increased mitochondrial NADH/NAD^+^, reductive mitochondrial stress, and ROS generation^52, 53^. The ability of lactate accumulation to stimulate mitochondrial ATP production also provided cells with an increased ability to engage in translation in support of cellular differentiation and effector function as shown for effector T cells.

Recently, roles of lactate acting as a cytosolic messenger to regulate ER magnesium homeostasis and cellular mitosis have been proposed^54, 55^. These functions of lactate are stereospecific for L-lactate. In contrast, lactate’s effects on ETC activity are not stereospecific. Both D- and L-lactate are effective at stimulating mitochondrial ATP production. This difference in stereospecificity may be due to the evolutionary role of the ETC. In prokaryotes, both D-lactate and L-lactate contribute to ETC activity^56^. Further studies are needed to define the molecular details of how lactate activates the ETC. However, the ability to significantly improve proliferation of antimycin treated cells and to overcome cytochrome B deficiency suggests a potential role in enhancing complex III electron transfer.

### The ability of D-lactate to act as a mitochondrial messenger offers translational opportunities

The general principles uncovered here also enabled us to study other forms of mitochondrial dysfunction such as chronically antigen-stimulated T cells and cells carrying mitochondrial DNA mutations. The ability of D-lactate to stimulate ETC activity to support sufficient *de novo* pyrimidine production for survival and growth of cells carrying mitochondrial disease mutations suggests it may be beneficial in the study of patients with inborn errors of metabolism or with acquired mitochondrial diseases. Treatments of mitochondrial diseases using metabolites that indirectly increase cellular reductive stress have led to significant side effects^56^. D-lactate has the benefit of stimulating ETC activity without creating reductive stress. D-lactate has been used in the treatments of wounds and critically ill patients for nearly 100 years as a component of lactated-Ringer’s solution, which contains a 28 mM racemic mix of L- and D-lactate^57^. Solutions containing up to 42 mM D-lactate have been shown to be safe in humans^58^.

Finally, the success of cellular immunotherapy approaches for cancer have been limited both *in vitro* and *in vivo* by T cell exhaustion associated with impaired mitochondrial function^59^. While L-lactate is often elevated in tumors, its metabolism generates NADH and has been associated with suppression of the immune response partly through redox stress^34^. In contrast, D-lactate, which bypasses cytosolic LDH metabolism while enhancing mitochondrial oxidative phosphorylation could have benefits for both *in vitro* grown T cells or T cell expansion *in vivo.* Skewing T cells away from glycolysis towards oxidative phosphorylation has been associated with improved anti-tumor effector responses^60^.

The ability to reduce mitochondrial reductive stress with D-lactate also has important potential applications for the study of other diseases associated with mitochondrial reductive stress including neurodegenerative diseases, cardiovascular disease, and aging. The ability to stimulate oxidative phosphorylation while reducing mitochondrial reductive stress provides a new tool to study the association of mitochondrial stress with a wide variety of disorders. Collectively, these results establish lactate as a major determinant of cellular ATP production and a critical regulator of the ability of oxidative phosphorylation to suppress glucose fermentation.

### Limitations of the Study

While our studies indicate lactate stimulates ETC activity independent of its metabolism, the precise molecular mechanism remains to be defined and will likely require detailed biochemical and structural studies in the future. Recent imaging probe have shown mitochondrial lactate levels to reach the millimolar range, which is significantly higher than that measured in the cytosol^27^, suggesting the mitochondria have evolved to retain and perhaps require high levels of lactate. Furthermore, given the prevalent use of D-lactate as a component of Ringer’s Lactate solution in humans, the in vivo implications of our findings require detailed characterization. As D-lactate is primarily produced by commensal bacteria and have been reported to reach up to 10mM in the portal vein^61^, its role in regulating gut and liver physiology also requires additional study.

## ACKNOWLEDGEMENTS

We thank members of the Thompson laboratory for critical discussions and Drs. Siqi Liu, Lydia Finley, and Ralph DeBerardinis for manuscript feedback. X.C. is supported by the NCI (K99 CA256505). This work is supported by grants from the NCI (P30 CA008748).

## AUTHOR CONTRIBUTIONS

X.C. and C.B.T. conceived the study. X.C. performed most experiments and analyzed the data, with assistance from O.J. C.N. and D.L. performed the T-cell experiments. T.S.F. and K.R. assisted in methods development. X.C. C.N. and C.B.T. interpreted the results and wrote the manuscript. All authors participated in discussing and finalizing the manuscript.

## DECALARATION OF INTERESTS

C.B.T. is a founder of Agios Pharmaceuticals. He is on the Board of Directors of Regeneron and Charles River Laboratories. The other authors declare that they have no competing interests.

---

**Supplemental Figure 1.**
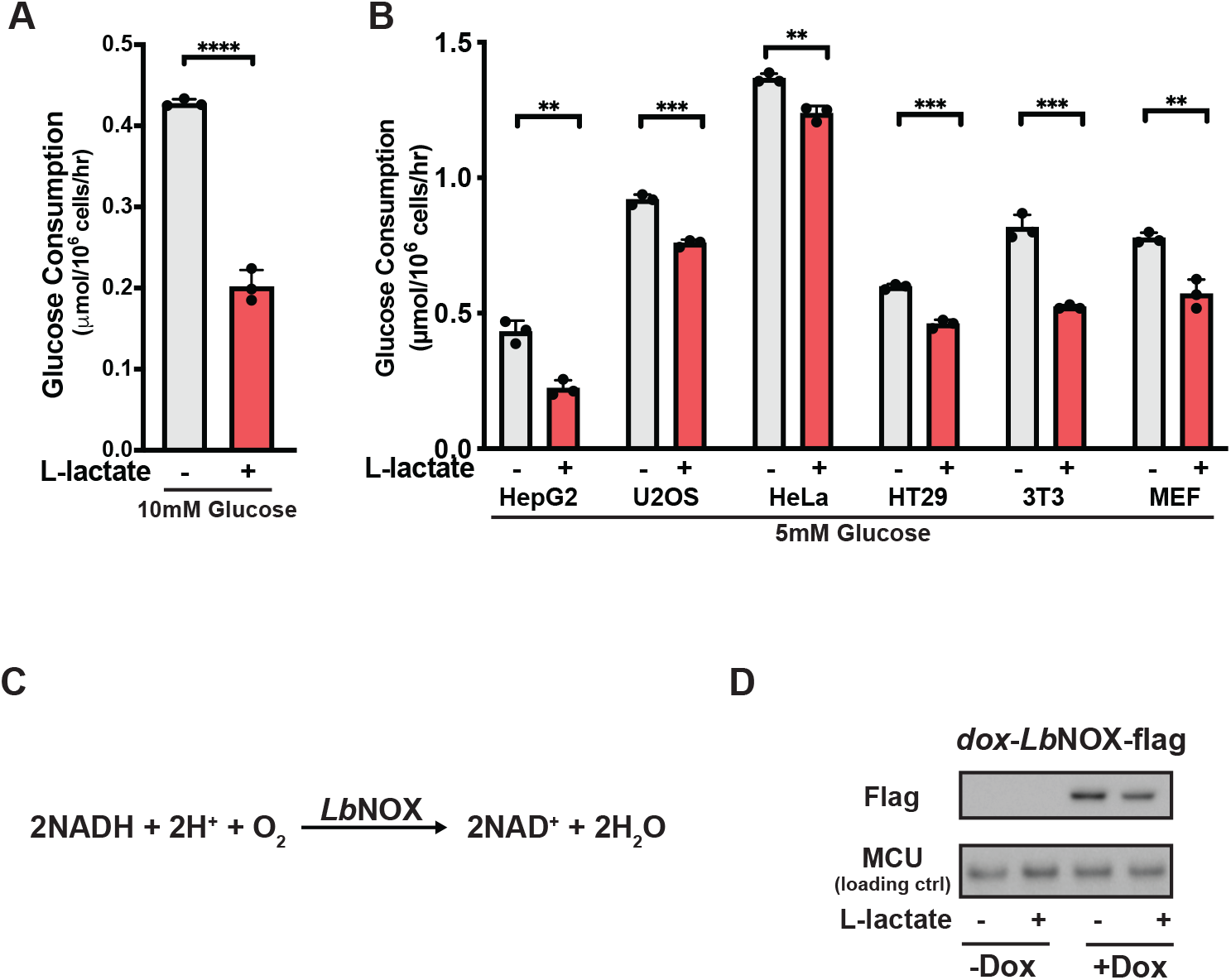
Supporting data for Figure 1. (A) Glucose consumption of HepG2 cells cultured in medium containing 10mM glucose with or without the addition of 20mM L-lactate. (B) Glucose consumption of indicated cell lines cultured in medium containing 5mM glucose with or without the addition of 10mM L-lactate. (C) Schematic of *Lb*NOX reaction. (D) Western blot of doxycycline-inducible *Lb*NOX-flag expression using a FLAG antibody. All error bars represent mean + SD with a minimum n of 3. Statistical analysis in (A) and (B) was performed using two-sided Student’s t-tests. **p < 0.01, ***p < 0.001, ****p < 0.0001.

**Supplemental Figure 2.**
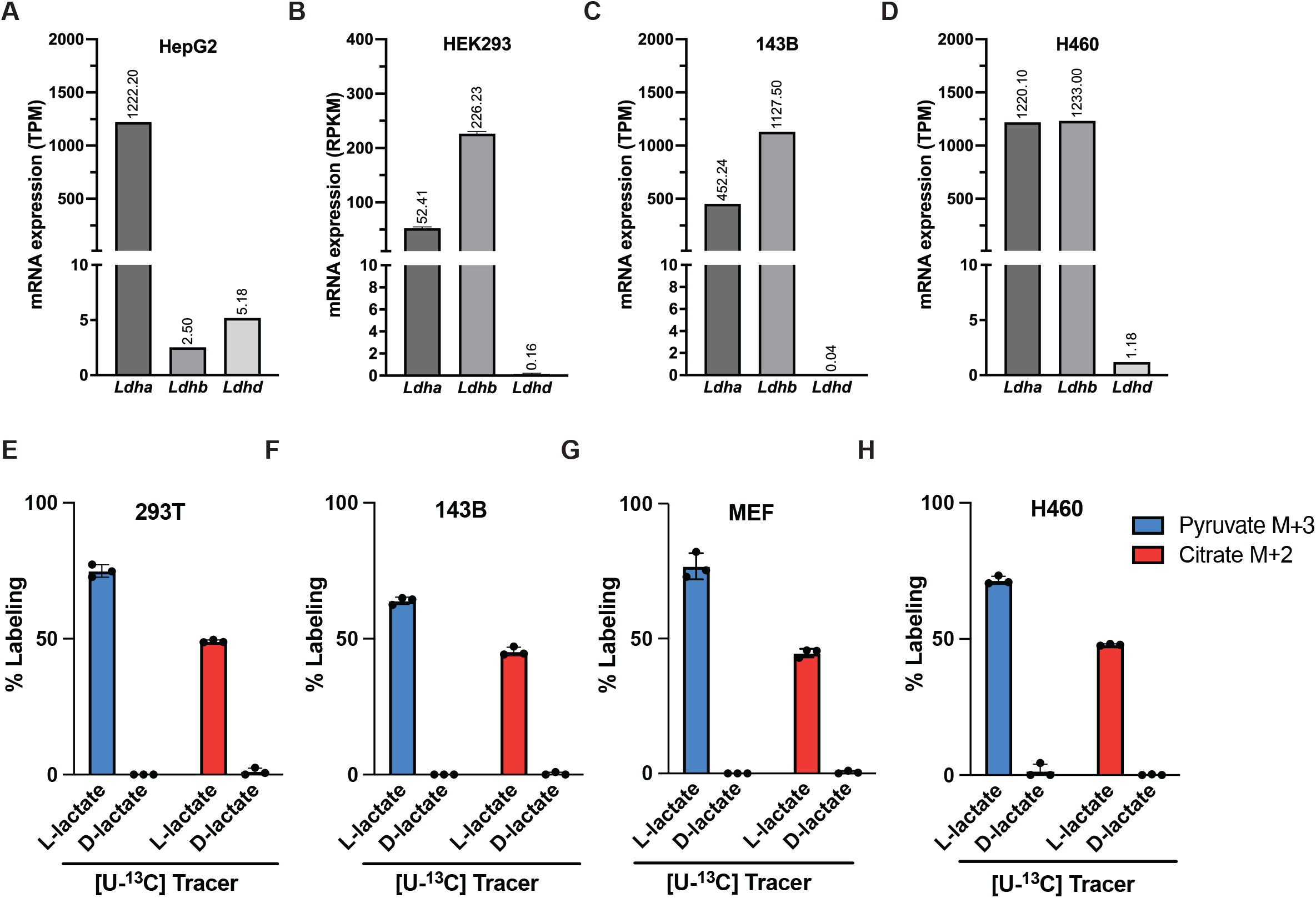
Supporting data for Figure 2. (A-D) mRNA expression of indicated LDH isoform from available RNA sequencing datasets. (A), (C), and (D) are derived from the CCLE dataset ref.^62^. (B) is derived from data from ref.^63^. TPM, transcript per million; RPKM, reads per kilobase per million mapped reads. (E-H) Percent labeling of pyruvate M+3 or citrate M+2 following incubation with 10mM [U-^13^C] L- or D-lactate for 8 hours in the indicated cell lines.

**Supplemental Figure 3.**
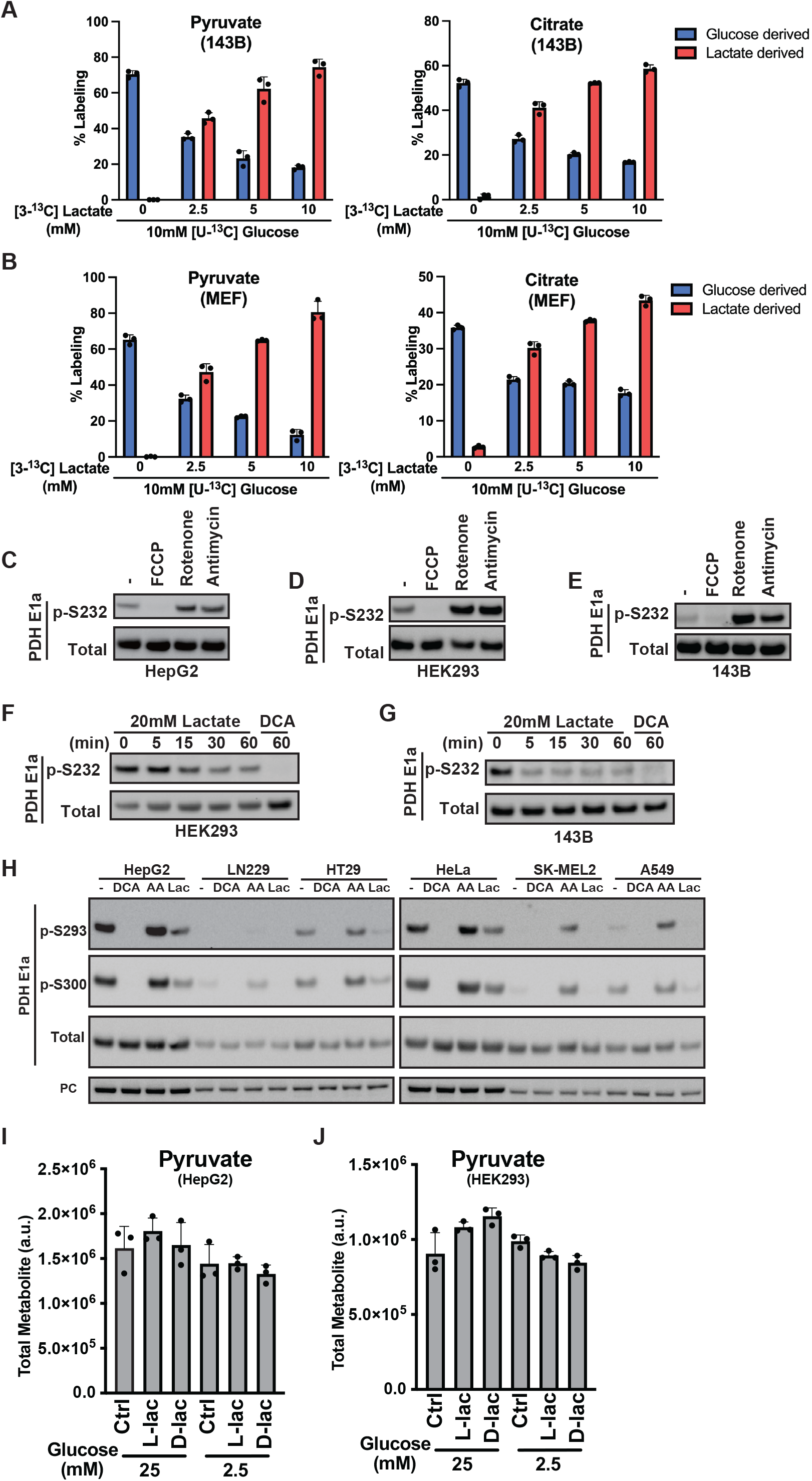
Supporting data for Figure 3. (A and B) Percent labeling of pyruvate and citrate from 10mM [U-^13^C] glucose and indicated concentration of [3-^13^C] lactate after 30 min incubation in (A) 143B and (B) MEF cells. (C) HepG2, (D) HEK293, and (E) 143B cells were treated with the indicated drugs (1µM) for 30 mins followed by immunoblotting analysis with the indicated antibodies. (F) HEK293 and (G) 143B cells were treated with the 20mM L-lactate or 5mM DCA for the indicated times followed by immunoblotting analysis with the indicated antibodies. (H) The indicated cell lines were treated with 5mM DCA, 1µM antimycin A, or 20mM L-lactate followed by immunoblotting analysis. (I) HepG2 and (J) HEK293 cells cultured in 25mM or 2.5mM glucose were incubated with 20mM NaCl, L- or D-lactate for 1 hour before total citrate abundance was analyzed using GC-MS. All error bars represent mean + SD with a minimum n of 3.

**Supplemental Figure 4.**
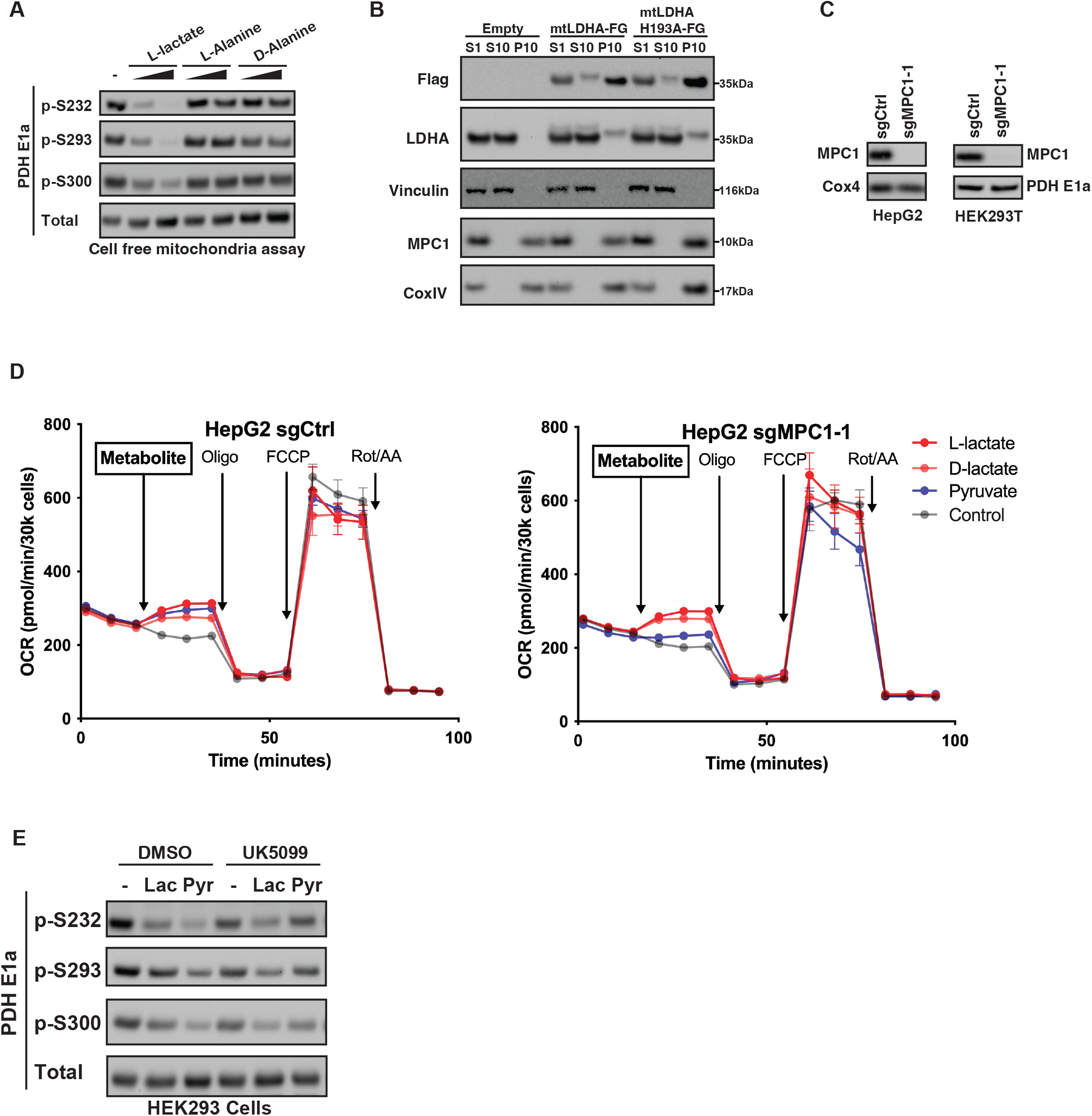
Supporting data for Figure 4. (A) Mitochondria purified from 293T cells were incubated in assay buffer with the indicated treatments (5mM and 15mM) for 30 mins at 30°C, followed by immunoblotting with the indicated antibodies. (B) Subcellular fractionation and immunoblotting analysis of 293T cells expressing the indicated mitochondrial-matrix (mt) targeted proteins with C-terminal Flag (FG) tag. S1, cytosol; P10, mitochondria fraction; S10, S1 minus P10. (C) Immunoblotting of cell populations transduced with the indicated CRISPR guides. (D) OCR of HepG2 cells containing the indicated sgRNA measured using Seahorse Bioanalyzer. Metabolite arrow indicates injection of 20mM NaCl, L- or D-lactate, or 2mM pyruvate. (E) Following 3 hour treatment with 50µM UK5099, cells were treated with 20mM NaCl, 20mM L-lactate, or 2mM pyruvate for 30 mins followed by immunoblotting analysis with the indicated antibodies. All error bars represent mean ± SD with a minimum n of 3.

**Supplemental Figure 5.**
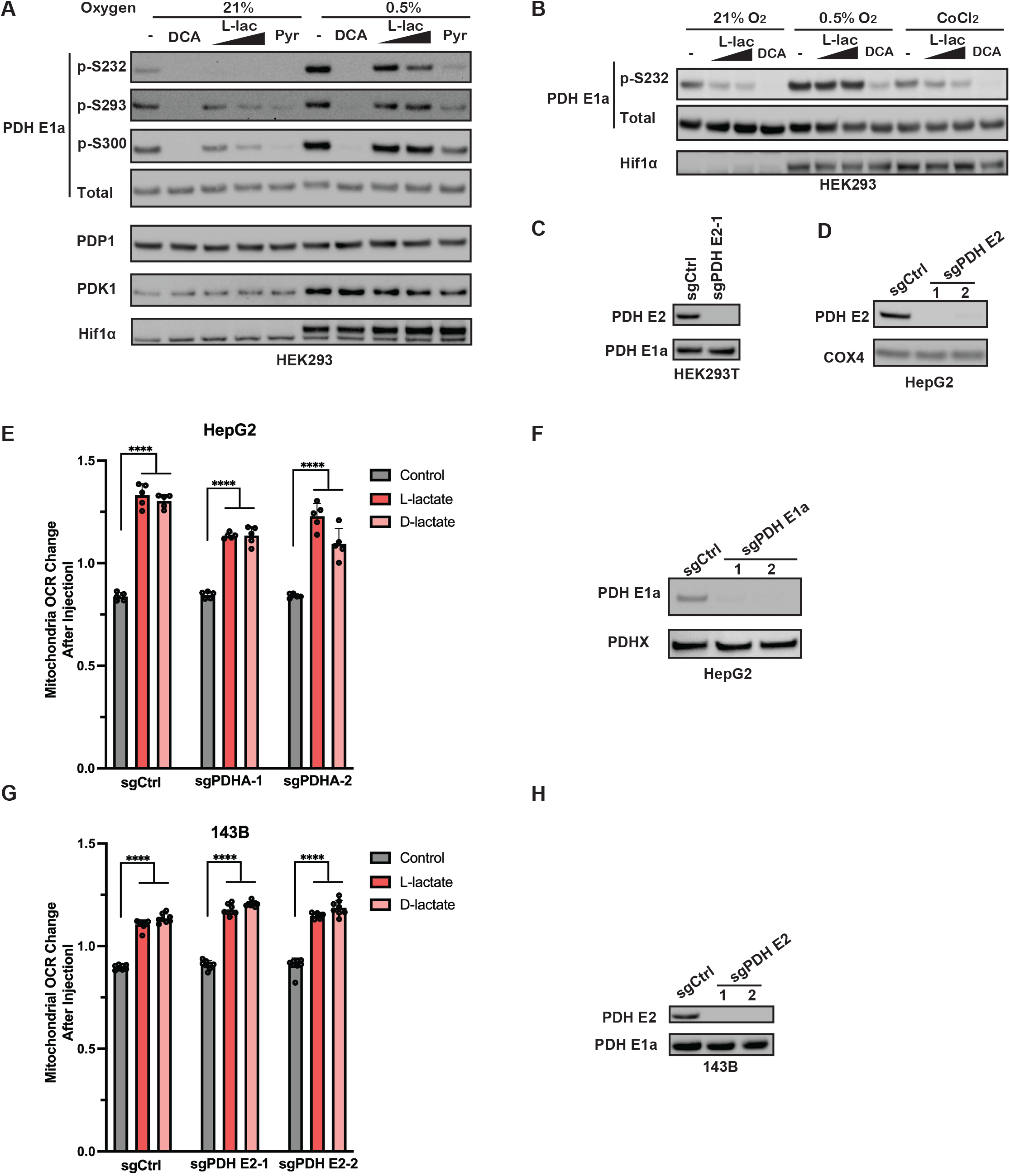
Supporting data for Figure 5. (A) Following overnight culture at the indicated oxygen concentration, HEK293 cells were treated with 5mM DCA, 15mM or 30mM L-lactate, or 2mM pyruvate for 30 mins at the indicated oxygen concentration, followed by immunoblotting. (B) Following overnight culture at the indicated oxygen concentration or in 100µM CoCl2, HEK293 cells were treated with 15mM or 30mM lactate or 5mM DCA for 30 mins followed by immunoblotting. (C and D) Immunoblotting of cell populations transduced with the indicated CRISPR guides. (E and G) OCR of HepG2 (E) or 143B (G) cells transduced with the indicated sgRNA measured using Seahorse Bioanalyzer and reported as the mitochondrial OCR change following injection of indicated metabolites. (F and H) Immunoblotting of cell populations transduced with the indicated CRISPR guides. All error bars in represent mean+SD with a minimum n of 3. Statistical analysis in (E) and (G) was performed using one way ANOVA. ****P<0.0001.

**Supplemental Figure 6.**
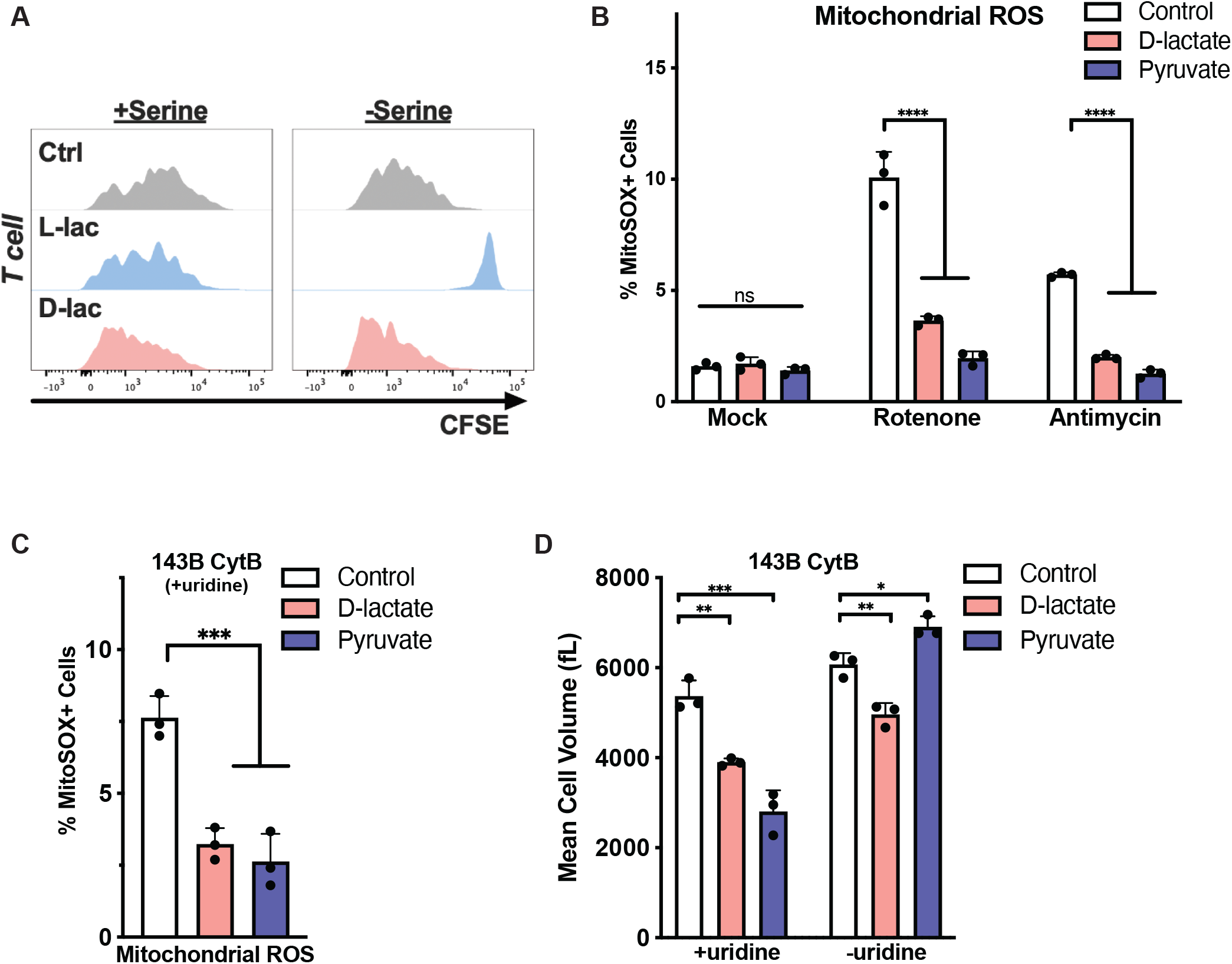
Supporting data for Figure 6. (A) CFSE dilution of CD4 T cells cultured in serine-deficient RMPI supplemented with NaCl (20mM), L-lactate (20mM), or D-lactate (20mM). (B) 143B cells were treated with 100nM of rotenone or antimycin A in the presence of NaCl (20mM), D-lactate (20mM) or pyruvate (2mM) for 30 minutes. Percentage of MitoSOX+ cells was then analyzed using FACS. (C) Percentage of MitoSOX+ cells following 6 days of culture in the indicated treatment conditions. (D) Mean cell volume of 143B CytB cells cultured in the indicated conditions for eight days was measured using the Multisizer 3. All error bars in represent mean+SD with a minimum n of 3. Statistical analysis in (B) and (C) was performed using one way ANOVA, in (D) was performed using two-sided Student’s t-test. *p < 0.05, **p < 0.01, *** p < 0.001, ****P<0.0001.

**Supplemental Figure 7.**
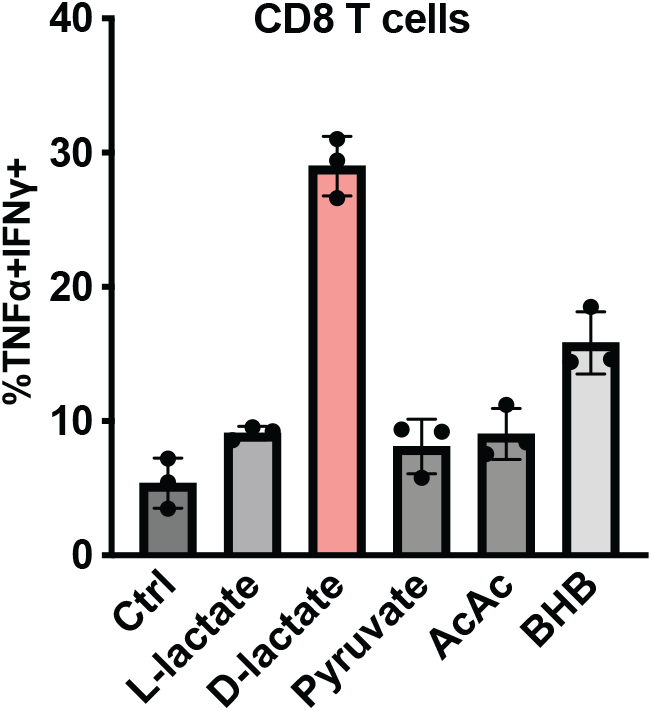
Supporting data for Figure 7. Percent of IFNγ and TNFα positive CD8 T cells cultured in complete RPMI with the indicated supplements. Ctrl, control; AcAc, acetoacetate; BHB, beta-hydroxybutyrate.

## EXPERIMENTAL MODEL AND SUBJECT DETAILS

### Cell lines

The 293T and HEK293 cell lines, the cancer cell lines HepG2, 143B, H460, HT29, HeLa, U2OS, A549, LN229, SK-MEL2, and the NIH 3T3 cell lines were obtained from the American Type Culture Collection (ATCC). The MEF cell line was derived by SV40 large T antigen immortalization. The 143B CytB cells were a gift from Dr. Ralph J. DeBerardinis at UT Southwestern and was previous established by Rana et. al^32^. The 143B Rho0 and matched wild type cells were obtained from Kerafast (Cat. ESA106). All cell lines were cultured in DMEM High Glucose (DMEM HG prepared by the MSKCC Media Core) supplemented with 10% FBS (Gemini), 100 unit/ml penicillin and 100 µg/ml streptomycin. The 143B CytB, 143B Rho0 and matched WT controls were also supplemented with 1mM sodium pyruvate and 250µM uridine. All cell lines were cultured in a 37°C incubator with 5% CO2. All cell lines were routinely verified to be mycoplasma-free by MycoAlert Mycoplasma Detection Kit (Lonza).

### Primary cell cultures

Cells were isolated from mouse spleen and lymph nodes and purified by negative selection with Dynabeads Untouched Mouse CD8 Cells Kit per manufacturer’s instructions (ThermoFisher). Purified CD8 T cells were activated as previously described^64^. Briefly, cells were seeded at 50,000 cells/well in 96-well plates precoated in 1mg/ml goat anti-hamster IgG (MP Biomedicals) at 1:20 dilution overnight at 4°C or 2-4h at 37°C. T cells were cultured for approximately 72h in standard RPMI (Fisher) or RPMI with 1mM glucose supplemented with 10% dialyzed FBS, 4 mM glutamine, 50 μM β-mercaptoethanol, 0.5μg/ml anti-CD3 (Fisher), 1µg/mL anti-CD28 (Fisher) and 100U/ml human IL-2 (Peprotech).

### Mice

C57BL/6J and IFN-γ reporter mice strain #017581 were purchased from Jackson Laboratories. All animal experiments described adhered to policies and practices approved by Memorial Sloan Kettering Cancer Center’s Institutional Animal Care and Use Committee (IACUC) and were conducted as per NIH guidelines for animal welfare (Protocol Number 11-03-007, Animal Welfare Assurance Number FW00004998). Experiments were performed using male and female mice aged 6–12 wk.

## METHODS DETAILS

### Cell proliferation

For cell proliferation experiments, cells were seeded in standard culture media described above. The following day, cells were washed with warm PBS x 2, followed by media change to the experimental tissue culture medium supplemented with dialyzed FBS (dFBS, Gemini). Unless otherwise stated, all proliferation experiments are done in High Glucose (25mM) DMEM based media with 4mM glutamine and without pyruvate. Galactose was used at a final concentration of

12.5mM. For HepG2 cell proliferation time course, cells were seeded at a density of 150k cells/well in 12 well plates and cultured in 1ml media with media change every 2 days. For galactose proliferation, 143B and MEFs were seeded at a density of 15k cells per well in 12 well plates, cultured in 2 ml of media with media change every 2 days. 143B CytB and Rho0 cells were seeded at 20k/well in a 12 well plate, cultured in 1ml of media with media change every 2 days. For ETC inhibitor proliferation, 143B and MEFs were seeded at a density of 7,000 and 10,000 cells per well, respectively, in 6 well plates and cultured in 2ml of media. Unless otherwise indicated, the following concentration of metabolites were added to standard DMEM

HG: 20mM NaCl, 20mM L-lactate, or 20mM D-lactate. Cells were counted using a Multisizer 3 Coulter Counter (Beckman) at indicated time points.

### PDH activation assay in cells

For PDH activation assays, cells were seeded in 6-well plates overnight. The next morning, media was refreshed with DMEM HG supplemented with 10% dFBS to enable cells to adapt to fresh serum and media. 60-90 mins later, media was changed to the indicated treatment (i.e. lactate, DCA, etc.) in DMEM HG + 10% dFBS for 30 minutes (or the indicated times during the time course) before cells were harvested for standard western blot analysis. 143B CytB and 143B Rho0 cells were seeded in media supplemented with pyruvate and uridine. The next morning, media change and treatments are done in DMEM HG + 10% dFBS. For hypoxia treatment, cells were plated in 6-well plates in either in normoxia (21% O2) or the hypoxia chamber (set to 0.5% O2) overnight. The next morning, media was changed to fresh DMEM + dFBS that had been equilibrated in standard incubator or the hypoxia chamber for at least one hour. 60-90 minutes later, media was changed to the indicated treatment (i.e. lactate, DCA) in equilibrated DMEM HG + 10% dFBS for 30 minutes before cells were harvested for standard western blot analysis.

### Subcellular fractionation and analysis of mitochondrial fractions

Mitochondria purification was carried out using standard subcellular fractionation protocols. Briefly, confluent 15cm dishes of 293T cells was washed and harvested in ice-cold buffer A (250 mM sucrose, 10mM KCl, 1.5mM MgCl2, 0.5 mM EGTA, 10mM Tris-HCl, pH 7.4), followed by douncing with a glass homogenizer over 25 strokes, and centrifuged at 1000g x 2 to remove the nuclei fraction and cell debris (P1). The remaining cytosolic fraction (S1) was centrifuged at 10,000g, and the pellet (P10) containing the mitochondria fraction was washed 2x in Buffer A before being resuspended in Buffer A containing additional 10mM KH2PO4 pH 7.4, 1mM Glutamate, 1mM Malate, and 0.5mM ADP (Buffer B). For *in vitro* PDH activity assay, 50µg of mitochondria was incubated with the appropriate treatment (i.e. DCA, lactate, pyruvate) resuspended in Buffer B in a final reaction volume of 20µl and incubated at 30°C for 30 mins. Unless otherwise indicated, the following concentrations were used: NaCl/DCA/L-lac/D-lac/Pyr/Calcium, 5mM; ATP/ADP, 4mM, Phosphatase Inhibitor: 1x. Rapid freeze and thaw was repeated 5x using liquid nitrogen and a 30°C water bath. The entire reaction mixture was then subjected to SDS-PAGE western blotting according to standard protocol described below.

For *in vitro* mitochondria TCA cycle tracing, cells were cultured in doxycycline containing media overnight to induce the expression of mito-LDHA WT or mito-LDH H193A or empty vector. The next day, cells were harvested and mitochondria fractions were obtained as described above. Then 100µg of mitochondria fraction was resuspended in Buffer A containing 10mM KH2PO4 pH 7.4, 1mM Malate, and 0.5mM ADP (Buffer C) along with the indicated treatments and [U-^13^C] metabolites. Following 30 min of incubation at 30°C, the entire reaction was quenched by adding 800µl of 80:20 methanol:water (kept in −80°C), followed by standard GC-MS analysis as described below.

NAD^+^/NADH measurements were done following manufacturer’s protocol with the modifications below (Biovision, K337). For mitochondrial NAD(H) measurements, 150µg of mitochondrial fraction was incubated with the appropriate treatment in 20µl of Buffer B at 30°C for 30 mins. The reaction was then quenched with the addition of 200ul ice-cold NADH/NAD extraction buffer, from which 100ul was immediately taken out and heated to 75°C for 30 mins to decompose NAD and the remaining samples were left on ice for 30 mins. Next, 50µl of each half was then transferred to a 96 well clear-bottom plate. Each assay plate also included wells that contained NADH standards (0, 20, 40, 60, 80, 100pmol) and control wells that contained the treatment metabolites/drug only (i.e. L-/D-lactate, FCCP, Rotenone) in extraction buffer to ensure no cross-reactivity with the enzymes used in the cycling assay. For each well, 98µl of NAD cycling buffer and 2µl of NAD cycling enzyme (100µl total) were added and incubated at room temperature for 5 mins to convert NAD^+^ to NADH. 10µl of NADH developer was then added to each well and allowed to incubate for up to 120 mins before sample OD at 450nm was read using a plate reader. The NAD^+^/NADH ratios were calculated according to manufacturer’s protocol by subtracting NADH (samples at 75°C) from total NAD (samples in ice) and dividing by NADH levels.

### Glucose consumption measurement

Cells were seeded at a density of 500k/well in 12 well plates. The next day, cells were washed with PBS x 2 and cultured in glucose-deficient DMEM (MSKCC media core) supplemented with the indicated final glucose concentrations and treatments. Starting media was incubated identically without cells. Following incubation for 6-8 hours, media glucose consumption was then measured using the YSI 2900 analyzer or GlucCell meter and normalized by cell number.

Media glucose concentration of ∼5mM was chosen after assessing the linear range of the media glucose analyzers and to minimize interference due to L-lactate produced from significant glycolysis.

### Gene knockout and overexpression

CRISPR-Cas9 mediated gene knockout was achieved using the lentiCRISPR v2 system (Addgene 52961 and 98292), and polyclonal cell populations were used for the experiments.

The guide sequences are indicated below. Mitochondrial-matrix targeted WT and H193A LDHA with a C-terminal FLAG tag were synthesized using gBlock (IDT) using the mitochondrial targeting sequence of subunit IV of human cytochrome c oxidase. Ectopic expression of mito-LDHA and *Lb*NOX (Addgene, 75285) was then achieved using the pINDUCER20 (Addgene, 44012) tet-on viral expression system. Doxycycline was used at 100ng/mL for gene induction overnight. Complete antibiotic selection was applied to all genetically modified cells before proceeding to experiments.

### Western blot

Cells were lysed in RIPA lysis buffer (Millipore 20-188) supplemented with protease and phosphatase inhibitors (Thermo Fisher, 78425 and 78428). Protein concentration was determined by BCA protein assay (Thermo Fisher, 23228), following which equal amount of protein was loaded and separated in polyacrylamide gels. Protein was then transferred to nitrocellulose membrane for immunoblotting.

### Metabolite analysis using GC-MS

All cells were seeded in 6-well plates overnight prior to metabolic analysis the following day at a density of 500k cells/well. For total metabolite analysis, media was changed to glucose deficient DMEM supplemented with the indicated glucose, 10% dialyzed FBS (Gemini), and indicated treatments prior to harvesting at the indicated time points in the figure legends. For [U-^13^C] L-lactate and [U-^13^C] pyruvate tracingin HEK293T cells, sgCtrl and sgMPC1 KO cells expressing mito-LDHA WT, H193A, or empty vector were cultured in 100ng/mL of Doxycycline overnight. The next day, media was refreshed with [U-^13^C]-L-lactate or [U-^13^C]-pyruvate for 3 hours. For [U-^13^C]-glucose and [3-^13^C]-lactate tracing experiments, cells were incubated with tracing media (DMEM+10% dFBS) for 30 minutes to minimize [3-^13^C]-lactate contribution to the second turn of the TCA cycle. The [U-^13^C] L-lactate and [U-^13^C] D-lactate tracing experiments were carried out over 8 hours in media containing 10mM of either tracer. Metabolism was quenched by the addition of 1 mL of 80:20 methanol:water and stored at −80°C overnight. The methanol-extracted metabolites were cleared by centrifugation and supernatant was dried in a vacuum evaporator (Genevac EZ-2 Elite) for 5 hours. Dried metabolites were dissolved in 40 mg/mL methoxyamine hydrochloride (Sigma, 226904) in pyridine (Thermo Fisher, TS-27530) for 90 min at 30°C and derivatized with MSTFA with 1% TMCS (Thermo Fisher, TS-48915) for 30 min at 37°C. Samples were analyzed using an Agilent 7890A GC connected to an Agilent 5975C Mass Selective Detector with electron impact ionization. The GC was operated in splitless mode with constant helium gas flow at 1 mL/min. 1 μL of derivatized metabolites was injected onto an HP-5MS column, the inlet temperature was 250°C, and the GC 6 oven temperature was ramped from 60 to 290°C over 25 min. Peak ion chromatograms for metabolites of interest were recorded and extracted at their specific m/z with MassHunter Quantitative Analysis software v10.0 (Agilent Technologies). Ions used for quantification of metabolite levels are as follows: α-ketoglutarate m/z 304; citrate m/z 465; lactate m/z 219; pyruvate m/z 174. All peaks were manually inspected and verified relative to known spectra for each metabolite. Natural isotope abundance correction was performed using IsoCor (https://isocor.readthedocs.io/en/latest/index.html).

### Oxygen consumption, extracellular acidification, and glycolytic proton efflux rate measurements

Oxygen consumption rate (OCR), extracellular acidification rate (ECAR), and glycolytic proton efflux rate (GlycoPER) were measured using a XFe96 Extracellular Flux Analyzer (Agilent) following manufacturer’s instructions. Cells were plated in Seahorse microplates (Agilent) at appropriate densities (10,000 cells/well for 143B or 30,000 cells/well for HepG2), and were allowed to adhere overnight. Cell culture media were then removed and replaced with Seahorse media (DMEM, Agilent 103575, supplemented with 10 mM glucose and 4 mM glutamine), unless stated otherwise in figure legends. OCR analysis was performed at basal level and after subsequent injections of oligomycin (2 μM), FCCP (0.5 μM), and rotenone plus antimycin A mix (each at 2 μM) according to the manufacturer’s instructions, unless stated otherwise in the figure legend. Immediately after measurements, cell number were determined using a Multisizer 3 Coulter Counter (Beckman). Results were then normalized to cell number. Mitochondrial OCR was determined by subtracting non-mitochondrial OCR (measurements after rotenone/antimycin addition) from that of the basal OCR. For T cells, Seahorse microplate wells were precoated in Poly-D-Lysine (ThermoFisher A3890401) at 0.1mg/ml for 30 minutes at room temperature followed by two PBS washes and stored overnight at 4°C. On the day of the assay, T cells were seeded at 100,000 per well in Seahorse RPMI media supplemented with 10mM glucose, 4 mM glutamine and 50 μM β-mercaptoethanol +/-indicated metabolites and OCR analysis was then performed as above.

### ROS measurement

143B cells were incubated with 0.5µM MitoSOX Red (ThermoFisher M36008) for 15 minutes in complete media (DMEM HG + 10% dFBS) to allow dye loading followed by wash x 2 in dye-free complete media before they were subjected to treatment with 100nM of rotenone or antimycin A in the presence of different metabolites for 30 minutes at 37°C / 5% CO2. Cells were then harvested by scraping and DAPI was added (0.2ug/ml) prior to flow cytometry. 143B CytB cells were cultured in the indicated treatment conditions for 6 days with media change every 2 days. On the day of the assay, cells were incubated with 0.5μM MitoSOX Red and 0.2μg/ml DAPI for 30 mins at 37°C/5% CO2, washed 2x in complete media and harvested using a cell scraper prior to flow cytometry.

### T-cell experiments

For monocarboxylate treatments, 100,000 cells were seeded in 200 ul/well in 96-well plates that were pre-coated with a 1:20 dilution of goat anti-hamster IgG in PBS (50µg/ml). CD8 T cells were primed and polarized with anti-CD3ε (0.25 mg/mL), anti-CD28 (1 mg/mL), and human IL-2 (100U/ml) for 24 hours prior to monocarboxylate treatment. Monocarboxylates were added at 20mM and cells were incubated for an additional 48 hours in 37°C/5% CO2 prior to analysis.

For serine deficient growth, Mouse CD4+ T cells were purified from lymph nodes and spleens of six to eight-week-old C57BL/6J mice using Invitrogen™ Dynabeads™ Untouched™ Mouse CD4 Cells Kit (ThermoFisher, 11415D). Following purification, cells were stained for CFSE (ThermoFisher, C34570) per manufacturer’s instructions for 5 minutes in PBS at room temperature. Cells were subsequently activated as above with RPMI deficient for serine using 10% heat-inactivated dialyzed FBS supplemented with 10U/ml penicillin-streptomycin, 4 mM L-glutamine, 50 mM beta-mercaptoethanol and Na-L-lactate, Na-D-lactate or NaCl at 20mM. After 72h incubation in 37°C/5% CO2, T cells were analyzed for CFSE dilution by flow cytometry.

### Translation assay

To measure translation with puromycin incorporation, CD8 T cells were activated for 24h and treated with either NaCl (20mM) or D-lactate (20mM) for an additional 48h. Puromycin was added to the culture (10µg/ml) for 10 minutes in full media at 37°C/5% CO2 followed by surface staining and fixation/permeabilization as described below.

### Flow cytometry

For cytokine measurements in T cells treated with different monocarboxylates, cells were restimulated with eBioscience Cell Stimulation Cocktail 1:500 (Thermofisher, 00-4970-93) and GolgiStop 1:1000 (Fisher Scientific, BDB554724) for 4h in complete RPMI. Cells were then surface stained in PBS for anti-CD8ɑ (Biolegend, clone: 53–6.7), anti-CD25 (Biolegend, clone: PC61) and Live/Dead Fixable Blue (Fisher, L23105) for 15 min at room temperature prior to fixation. Following surface staining, cells were fixed/permeabilized (BD Biosciences) according to the manufacturer’s protocol. Intracellular staining was performed for anti-TNFα (Biolegend, clone: MP6-XT22) and anti-IFNγ (Biolegend, clone: XMG1.2) for 30 minutes at room temperature. For IFN-g reporter CD8 T cells grown in low glucose, T cells were harvested from 96-well plates and stained in PBS with anti-CD8ɑ 1:400 (Biolegend, clone: 53–6.7) and DAPI (0.2µg/ml) for 20 minutes on ice followed by immediate flow cytometry analysis.Intracellular staining with anti-puromycin was performed at 1:200 for 30 minutes at room temperature (Biolegend, clone: 2A4). Flow cytometry was performed on an LSRFortessa (BD Biosciences), and analysis was performed using Flowjo.

## QUANTIFICATION AND STATISTICAL ANALYSIS

Statistical analysis was calculated using Prism software (GraphPad). All error bars represent standard deviation with a minimal n = 3. Experiments were independently replicated, and representative data are shown. Paired two-tailed Student’s t-tests were used to ascertain statistical significance between two groups, and one-way ANOVA was used to assess statistical significance between three or more groups with one experimental parameter; * p < 0.05, ** p< 0.01, *** p < 0.001, **** p < 0.0001, ns; not significant. See figure legends for more information on statistical tests.

## REFERENCES

1. Amaral, J.F., Shearer, J.D., Mastrofrancesco, B., Gann, D.S., and Caldwell, M.D. (1986). Can lactate be used as a fuel by wounded tissue? Surgery 100, 252–261.

2. Gallagher, C.N., Carpenter, K.L., Grice, P., Howe, D.J., Mason, A., Timofeev, I., Menon, D.K., Kirkpatrick, P.J., Pickard, J.D., Sutherland, G.R., and Hutchinson, P.J. (2009). The human brain utilizes lactate via the tricarboxylic acid cycle: a 13C-labelled microdialysis and high-resolution nuclear magnetic resonance study. Brain 132, 2839–2849. 10.1093/brain/awp202.

3. Levasseur, J.E., Alessandri, B., Reinert, M., Clausen, T., Zhou, Z., Altememi, N., and Bullock, M.R. (2006). Lactate, not glucose, up-regulates mitochondrial oxygen consumption both in sham and lateral fluid percussed rat brains. Neurosurgery 59, 1122–1130; discussion 1130-1121. 10.1227/01.Neu.0000245581.00908.Af.

4. Gladden, L.B. (2004). Lactate metabolism: a new paradigm for the third millennium. J Physiol 558, 5–30. 10.1113/jphysiol.2003.058701.

5. Hui, S., Ghergurovich, J.M., Morscher, R.J., Jang, C., Teng, X., Lu, W., Esparza, L.A., Reya, T., Le Zhan, L., Yanxiang Guo, J., et al. (2017). Glucose feeds the TCA cycle via circulating lactate. Nature 551, 115–118. 10.1038/nature24057.

6. Faubert, B., Li, K.Y., Cai, L., Hensley, C.T., Kim, J., Zacharias, L.G., Yang, C., Do, Q.N., Doucette, S., Burguete, D., et al. (2017). Lactate Metabolism in Human Lung Tumors. Cell 171, 358–371.e359. 10.1016/J.CELL.2017.09.019.

7. Jang, C., Hui, S., Zeng, X., Cowan, A.J., Wang, L., Chen, L., Morscher, R.J., Reyes, J., Frezza, C., Hwang, H.Y., et al. (2019). Metabolite Exchange between Mammalian Organs Quantified in Pigs. Cell Metabolism 30, 594–606.e593. 10.1016/j.cmet.2019.06.002.

8. Bartman, C.R., Weilandt, D.R., Shen, Y., Lee, W.D., Han, Y., TeSlaa, T., Jankowski, C.S.R., Samarah, L., Park, N.R., da Silva-Diz, V., et al. (2023). Slow TCA flux and ATP production in primary solid tumours but not metastases. Nature 614, 349–357. 10.1038/s41586-022-05661-6.

9. Colegio, O.R., Chu, N.-Q., Szabo, A.L., Chu, T., Rhebergen, A.M., Jairam, V., Cyrus, N., Brokowski, C.E., Eisenbarth, S.C., Phillips, G.M., et al. (2014). Functional polarization of tumour-associated macrophages by tumour-derived lactic acid. Nature 513, 559–563. 10.1038/nature13490.

10. Carmona-Fontaine, C., Deforet, M., Akkari, L., Thompson, C.B., Joyce, J.A., and Xavier, J.B. (2017). Metabolic origins of spatial organization in the tumor microenvironment. Proceedings of the National Academy of Sciences of the United States of America 114, 2934–2939. 10.1073/pnas.1700600114.

11. Tasdogan, A., Faubert, B., Ramesh, V., Ubellacker, J.M., Shen, B., Solmonson, A., Murphy, M.M., Gu, Z., Gu, W., Martin, M., et al. (2020). Metabolic heterogeneity confers differences in melanoma metastatic potential. Nature 577, 115–120. 10.1038/s41586-019-1847-2.

12. Frauwirth, K.A., Riley, J.L., Harris, M.H., Parry, R.V., Rathmell, J.C., Plas, D.R., Elstrom, R.L., June, C.H., and Thompson, C.B. (2002). The CD28 signaling pathway regulates glucose metabolism. Immunity 16, 769–777. 10.1016/s1074-7613(02)00323-0.

13. Sullivan, Lucas B., Gui, Dan Y., Hosios, Aaron M., Bush, Lauren N., Freinkman, E., and Vander Heiden, Matthew G. (2015). Supporting Aspartate Biosynthesis Is an Essential Function of Respiration in Proliferating Cells. Cell 162, 552–563. 10.1016/J.CELL.2015.07.017.

14. Titov, D.V., Cracan, V., Goodman, R.P., Peng, J., Grabarek, Z., and Mootha, V.K. (2016). Complementation of mitochondrial electron transport chain by manipulation of the NAD+/NADH ratio. Science 352, 231–235. 10.1126/science.aad4017.

15. Pasteur, L. (1861). Expériences et vues nouvelles sur la nature des fermentations.

16. Racker, E. (1974). History of the Pasteur effect and its pathobiology. Molecular and cellular biochemistry 5, 17–23.

17. Kim, M.J., and Whitesides, G.M. (1988). L-Lactate dehydrogenase: substrate specificity and use as a catalyst in the synthesis of homochiral 2-hydroxy acids. Journal of the American Chemical Society 110, 2959–2964. 10.1021/ja00217a044.

18. Flick, M.J., and Konieczny, S.F. (2002). Identification of putative mammalian D-lactate dehydrogenase enzymes. Biochem Biophys Res Commun 295, 910–916. 10.1016/s0006-291x(02)00768-4.

19. de Bari, L., Atlante, A., Guaragnella, N., Principato, G., and Passarella, S. (2002). D-Lactate transport and metabolism in rat liver mitochondria. Biochem J 365, 391–403. 10.1042/bj20020139.

20. Patel, M.S., Nemeria, N.S., Furey, W., and Jordan, F. (2014). The pyruvate dehydrogenase complexes: structure-based function and regulation. J Biol Chem 289, 16615–16623. 10.1074/jbc.R114.563148.

21. Kato, M., Li, J., Chuang, J.L., and Chuang, D.T. (2007). Distinct structural mechanisms for inhibition of pyruvate dehydrogenase kinase isoforms by AZD7545, dichloroacetate, and radicicol. Structure 15, 992–1004. 10.1016/j.str.2007.07.001.

22. Liao, P.C., Bergamini, C., Fato, R., Pon, L.A., and Pallotti, F. (2020). Isolation of mitochondria from cells and tissues. Methods Cell Biol 155, 3–31. 10.1016/bs.mcb.2019.10.002.

23. Rogers, G.W., Brand, M.D., Petrosyan, S., Ashok, D., Elorza, A.A., Ferrick, D.A., and Murphy, A.N. (2011). High throughput microplate respiratory measurements using minimal quantities of isolated mitochondria. PLoS One 6, e21746. 10.1371/journal.pone.0021746.

24. Paxton, R., and Harris, R.A. (1984). Clofibric acid, phenylpyruvate, and dichloroacetate inhibition of branched-chain alpha-ketoacid dehydrogenase kinase in vitro and in perfused rat heart. Arch Biochem Biophys 231, 58–66. 10.1016/0003-9861(84)90362-x.

25. Halestrap, A.P. (2012). The monocarboxylate transporter family-Structure and functional characterization. IUBMB Life 64, 1–9. 10.1002/iub.573.

26. Chen, Y., Jr., Mahieu, N.G., Huang, X., Singh, M., Crawford, P.A., Johnson, S.L., Gross, R.W., Schaefer, J., and Patti, G.J. (2016). Lactate metabolism is associated with mammalian mitochondria. Nature Chemical Biology 12, 937–943. 10.1038/nchembio.2172.

27. Li, X., Zhang, Y., Xu, L., Wang, A., Zou, Y., Li, T., Huang, L., Chen, W., Liu, S., Jiang, K., et al. (2023). Ultrasensitive sensors reveal the spatiotemporal landscape of lactate metabolism in physiology and disease. Cell Metab 35, 200–211.e209. 10.1016/j.cmet.2022.10.002.

28. Xie, H., and Simon, M.C. (2017). Oxygen availability and metabolic reprogramming in cancer. J Biol Chem 292, 16825–16832. 10.1074/jbc.R117.799973.

29. Chandel, N.S., and Schumacker, P.T. (1999). Cells depleted of mitochondrial DNA (rho0) yield insight into physiological mechanisms. FEBS Lett 454, 173–176. 10.1016/s0014-5793(99)00783-8.

30. Spadafora, D., Kozhukhar, N., Chouljenko, V.N., Kousoulas, K.G., and Alexeyev, M.F. (2016). Methods for Efficient Elimination of Mitochondrial DNA from Cultured Cells. PLoS One 11, e0154684. 10.1371/journal.pone.0154684.

31. Diehl, F.F., Lewis, C.A., Fiske, B.P., and Vander Heiden, M.G. (2019). Cellular redox state constrains serine synthesis and nucleotide production to impact cell proliferation. Nat Metab 1, 861–867. 10.1038/s42255-019-0108-x.

32. Nikooie, R., Moflehi, D., and Zand, S. (2021). Lactate regulates autophagy through ROS-mediated activation of ERK1/2/m-TOR/p-70S6K pathway in skeletal muscle. J Cell Commun Signal 15, 107–123. 10.1007/s12079-020-00599-8.

33. Tauffenberger, A., Fiumelli, H., Almustafa, S., and Magistretti, P.J. (2019). Lactate and pyruvate promote oxidative stress resistance through hormetic ROS signaling. Cell Death Dis 10, 653. 10.1038/s41419-019-1877-6.

34. Quinn, W.J., 3rd, Jiao, J., TeSlaa, T., Stadanlick, J., Wang, Z., Wang, L., Akimova, T., Angelin, A., Schäfer, P.M., Cully, M.D., et al. (2020). Lactate Limits T Cell Proliferation via the NAD(H) Redox State. Cell Rep 33, 108500. 10.1016/j.celrep.2020.108500.

35. Robinson, B.H., Petrova-Benedict, R., Buncic, J.R., and Wallace, D.C. (1992). Nonviability of cells with oxidative defects in galactose medium: A screening test for affected patient fibroblasts. Biochemical Medicine and Metabolic Biology 48, 122–126. https://doi.org/10.1016/0885-4505(92)90056-5.

36. Bayona-Bafaluy, M.P., Sánchez-Cabo, F., Fernández-Silva, P., Pérez-Martos, A., and Enríquez, J.A. (2011). A genome-wide shRNA screen for new OxPhos related genes. Mitochondrion 11, 467–475. https://doi.org/10.1016/j.mito.2011.01.007.

37. Arroyo, J.D., Jourdain, A.A., Calvo, S.E., Ballarano, C.A., Doench, J.G., Root, D.E., and Mootha, V.K. (2016). A Genome-wide CRISPR Death Screen Identifies Genes Essential for Oxidative Phosphorylation. Cell Metabolism 24, 875–885. 10.1016/j.cmet.2016.08.017.

38. King, M.P., and Attardi, G. (1989). Human Cells Lacking mtDNA: Repopulation with Exogenous Mitochondria by Complementation. Science 246, 500–503. doi:10.1126/science.2814477.

39. Birsoy, K., Wang, T., Chen, W.W., Freinkman, E., Abu-Remaileh, M., and Sabatini, D.M. (2015). An Essential Role of the Mitochondrial Electron Transport Chain in Cell Proliferation Is to Enable Aspartate Synthesis. Cell. 10.1016/j.cell.2015.07.016.

40. Rana, M., de Coo, I., Diaz, F., Smeets, H., and Moraes, C.T. (2000). An out-of-frame cytochrome b gene deletion from a patient with parkinsonism is associated with impaired complex III assembly and an increase in free radical production. Ann Neurol 48, 774–781.

41. Weinberg, F., Hamanaka, R., Wheaton, W.W., Weinberg, S., Joseph, J., Lopez, M., Kalyanaraman, B., Mutlu, G.M., Budinger, G.R., and Chandel, N.S. (2010). Mitochondrial metabolism and ROS generation are essential for Kras-mediated tumorigenicity. Proc Natl Acad Sci U S A 107, 8788–8793. 10.1073/pnas.1003428107.

42. Bell, E.L., Klimova, T.A., Eisenbart, J., Moraes, C.T., Murphy, M.P., Budinger, G.R., and Chandel, N.S. (2007). The Qo site of the mitochondrial complex III is required for the transduction of hypoxic signaling via reactive oxygen species production. J Cell Biol 177, 1029–1036. 10.1083/jcb.200609074.

43. Themeli, M., Rivière, I., and Sadelain, M. (2015). New cell sources for T cell engineering and adoptive immunotherapy. Cell Stem Cell 16, 357–366. 10.1016/j.stem.2015.03.011.

44. Vardhana, S.A., Hwee, M.A., Berisa, M., Wells, D.K., Yost, K.E., King, B., Smith, M., Herrera, P.S., Chang, H.Y., Satpathy, A.T., et al. (2020). Impaired mitochondrial oxidative phosphorylation limits the self-renewal of T cells exposed to persistent antigen. Nat Immunol 21, 1022–1033. 10.1038/s41590-020-0725-2.

45. Yu, Y.-R., Imrichova, H., Wang, H., Chao, T., Xiao, Z., Gao, M., Rincon-Restrepo, M., Franco, F., Genolet, R., Cheng, W.-C., et al. (2020). Disturbed mitochondrial dynamics in CD8+ TILs reinforce T cell exhaustion. Nature Immunology 21, 1540–1551. 10.1038/s41590-020-0793-3.

46. Jones, R.G., Plas, D.R., Kubek, S., Buzzai, M., Mu, J., Xu, Y., Birnbaum, M.J., and Thompson, C.B. (2005). AMP-Activated Protein Kinase Induces a p53-Dependent Metabolic Checkpoint. Molecular Cell 18, 283–293. https://doi.org/10.1016/j.molcel.2005.03.027.

47. Chang, C.H., Curtis, J.D., Maggi, L.B., Jr., Faubert, B., Villarino, A.V., O’Sullivan, D., Huang, S.C., van der Windt, G.J., Blagih, J., Qiu, J., et al. (2013). Posttranscriptional control of T cell effector function by aerobic glycolysis. Cell 153, 1239–1251. 10.1016/j.cell.2013.05.016.

48. Rabinowitz, J.D., and Enerbäck, S. (2020). Lactate: the ugly duckling of energy metabolism. Nat Metab 2, 566–571. 10.1038/s42255-020-0243-4.

49. Im, M.J., and Hoopes, J.E. (1970). Energy metabolism in healing skin wounds. J Surg Res 10, 459–464. 10.1016/0022-4804(70)90070-3.

50. Goodwin, M.L., Harris, J.E., Hernández, A., and Gladden, L.B. (2007). Blood lactate measurements and analysis during exercise: a guide for clinicians. J Diabetes Sci Technol 1, 558–569. 10.1177/193229680700100414.

51. Andersen, L.W., Mackenhauer, J., Roberts, J.C., Berg, K.M., Cocchi, M. N., and Donnino, M.W. (2013). Etiology and therapeutic approach to elevated lactate levels. Mayo Clin Proc 88, 1127–1140. 10.1016/j.mayocp.2013.06.012.

52. Schwörer, S., Berisa, M., Violante, S., Qin, W., Zhu, J., Hendrickson, R.C., Cross, J.R., and Thompson, C.B. (2020). Proline biosynthesis is a vent for TGFβ-induced mitochondrial redox stress. Embo j 39, e103334. 10.15252/embj.2019103334.

53. Luengo, A., Li, Z., Gui, D.Y., Sullivan, L.B., Zagorulya, M., Do, B.T., Ferreira, R., Naamati, A., Ali, A., Lewis, C.A., et al. (2021). Increased demand for NAD(+) relative to ATP drives aerobic glycolysis. Mol Cell 81, 691–707.e696. 10.1016/j.molcel.2020.12.012.

54. Daw, C.C., Ramachandran, K., Enslow, B.T., Maity, S., Bursic, B., Novello, M.J., Rubannelsonkumar, C.S., Mashal, A.H., Ravichandran, J., Bakewell, T.M., et al. (2020). Lactate Elicits ER-Mitochondrial Mg(2+) Dynamics to Integrate Cellular Metabolism. Cell 183, 474–489.e417. 10.1016/j.cell.2020.08.049.

55. Liu, W., Wang, Y., Bozi, L.H.M., Fischer, P.D., Jedrychowski, M.P., Xiao, H., Wu, T., Darabedian, N., He, X., Mills, E.L., et al. (2023). Lactate regulates cell cycle by remodelling the anaphase promoting complex. Nature 616, 790–797. 10.1038/s41586-023-05939-3.

56. Ingledew, W.J., and Poole, R.K. (1984). The respiratory chains of Escherichia coli. Microbiol Rev 48, 222–271. 10.1128/mr.48.3.222-271.1984.

57. Semler, M.W., Self, W.H., Wanderer, J.P., Ehrenfeld, J.M., Wang, L., Byrne, D.W., Stollings, J.L., Kumar, A.B., Hughes, C.G., Hernandez, A., et al. (2018). Balanced Crystalloids versus Saline in Critically Ill Adults. New England Journal of Medicine 378, 829–839. 10.1056/NEJMoa1711584.

58. Kuze, S., Naruse, T., Yamazaki, M., Hirota, K., Ito, Y., and Miyahara, T. (1992). Effects of sodium L-lactate and sodium racemic lactate on intraoperative acid-base status. Anesth Analg 75, 702–707. 10.1213/00000539-199211000-00008.

59. Bader, J.E., Voss, K., and Rathmell, J.C. (2020). Targeting Metabolism to Improve the Tumor Microenvironment for Cancer Immunotherapy. Mol Cell 78, 1019–1033. 10.1016/j.molcel.2020.05.034.

60. Sukumar, M., Liu, J., Ji, Y., Subramanian, M., Crompton, J.G., Yu, Z., Roychoudhuri, R., Palmer, D.C., Muranski, P., Karoly, E.D., et al. (2013). Inhibiting glycolytic metabolism enhances CD8+ T cell memory and antitumor function. J Clin Invest 123, 4479–4488. 10.1172/jci69589.

61. McDonald, B., Zucoloto, A.Z., Yu, I.L., Burkhard, R., Brown, K., Geuking, M.B., and McCoy, K.D. (2020). Programing of an Intravascular Immune Firewall by the Gut Microbiota Protects against Pathogen Dissemination during Infection. Cell Host Microbe 28, 660–668.e664. 10.1016/j.chom.2020.07.014.

62. Ghandi, M., Huang, F.W., Jané-Valbuena, J., Kryukov, G.V., Lo, C.C., McDonald, E.R., Barretina, J., Gelfand, E.T., Bielski, C.M., Li, H., et al. (2019). Next-generation characterization of the Cancer Cell Line Encyclopedia. Nature 569, 503–508. 10.1038/s41586-019-1186-3.

63. Sun, Z., Xue, S., Xu, H., Hu, X., Chen, S., Yang, Z., Yang, Y., Ouyang, J., and Cui, H. (2019). Effects of NSUN2 deficiency on the mRNA 5-methylcytosine modification and gene expression profile in HEK293 cells. Epigenomics 11, 439–453. 10.2217/epi-2018-0169.

64. Ng, C., Aichinger, M., Nguyen, T., Au, C., Najar, T., Wu, L., Mesa, K.R., Liao, W., Quivy, J.P., Hubert, B., et al. (2019). The histone chaperone CAF-1 cooperates with the DNA methyltransferases to maintain Cd4 silencing in cytotoxic T cells. Genes Dev 33, 669–683. 10.1101/gad.322024.118.

